# Shifts in neural representations, not gaze, track how people reinterpret ambiguous social scenes

**DOI:** 10.64898/2026.10.06.749912

**Authors:** Clara A. Sava-Segal, Tory G. Benson, Emily S. Finn

**Affiliations:** Department of Psychological and Brain Sciences, Dartmouth College, Hanover, NH, USA 03755; Wu Tsai Institute, Yale University, New Haven, CT, USA, 06510; Department of Psychology, Yale University, New Haven, CT, USA, 06510; Department of Psychology, Rutgers University, New Brunswick, NJ, USA, 08901

## Abstract

How do we come to “see” the same thing differently after hearing someone else’s perspective? Revising an interpretation could reflect changes in how the brain represents the meaning of what we see, changes in how we visually sample the input, or both. To disentangle these mechanisms, we developed *MindLibs,* an adaptive fMRI and eye-tracking paradigm in which participants generate interpretations of ambiguous photographs and then view alternative interpretations chosen in real time to vary in semantic content from their own. Exposure to semantically distant alternatives revealed a strong link between reinterpretation as reported behaviorally and representational shifts in association cortex, but no systematic relationship between reinterpretation and changes in gaze. In particular, representational shifts in lateral default-mode network regions (DMN-B) predicted reinterpretation independently of shifts in frontoparietal, attention, and visual networks, and independently of whether gaze changed. In contrast, shifts in visual cortex tracked reinterpretation only when gaze also shifted, and contributed no unique variance beyond DMN-B. These findings identify higher-order conceptual reconfiguration, rather than perceptual resampling, as the primary account of how people revise the meaning of an ambiguous scene.

## Introduction

What makes someone change their mind? In everyday life, people regularly encounter perspectives that invite them to reinterpret something they thought they already understood: a friend’s take on a film they both watched, a therapist’s reframing of a difficult experience, or a critical review of a book they just enjoyed. Although people are often reluctant to revise an interpretation once it has formed ^1–3^, the interpretations offered by others can nevertheless sometimes prompt such revision ^4,5^. Yet it remains unclear what determines when reinterpretation occurs and how a stable sensory input comes to acquire a different meaning.

A useful perceptual analog of arriving at a new understanding of an unchanged stimulus comes from bistable perception, in which the same sensory input can give rise to alternating subjective percepts. Classic stimuli such as the Necker cube or the rabbit-duck illusion make it possible to dissociate changes in subjective experience from changes in sensory stimulation. Transitions between percepts in these paradigms are reflected in viewing behavior, with different percepts associated with distinct fixation patterns and oculomotor events ^6–8^. Further, these transitions have been linked to activity in both visual cortex and higher-order frontoparietal regions, suggesting that changes in what is seen involve interactions between sensory representations and systems that select among competing alternatives (e.g., ^9–12)^. Thus, shifts in subjective interpretation of an unchanged stimulus can coincide with changes in both visual sampling and neural representations, though the temporal order of these changes, and whether—and if so, which—one causes the other is debated ^13,14^.

Classic bistable stimuli typically involve two simple, mutually exclusive percepts, meaning “reinterpretation” is akin to switching between a fixed set of alternatives. Ambiguity in everyday life, on the other hand, is often richer and less resolvable: the same input can support multiple plausible meanings, with no single interpretation being objectively correct. In particular, naturalistic visual scenes containing social information can often be interpreted in many different ways. Reinterpreting more complex visual input may therefore require integrating new information into a graded understanding rather than simply choosing between a small number of discrete alternatives. This contrast motivates a distinction between two forms of updating that may support reinterpretation of a complex scene: a shift in the sensory input the visual system receives from the image (e.g., via gaze changes), and a shift in the meaning or category assigned to the image. We refer to the former as *visual resampling,* i.e., where and how the eyes move across the stimulus. We focus specifically on overt sampling as indexed by eye movements, rather than covert shifts in attention that can occur without any accompanying change in gaze ^15,16^. Existing work suggests that gaze behavior is shaped by the viewer’s task, goals, and current priorities rather than by properties of the image alone ^17–21^, raising the possibility that reinterpretation changes how an image is visually sampled as either a cause or a consequence of changing what is relevant or meaningful in the image. We refer to shifts in meaning or category as *conceptual updating*: understanding the image as depicting a different object, event, social situation, or intention. In principle, this can occur without any change in overt visual sampling, and should be expressed as a change in how the brain represents the same, unchanging visual input.

Characterizing how visual and conceptual updating operate during reinterpretation therefore requires a framework that can concurrently track changes in neural representations, changes in visual sampling, and changes in meaning, particularly when reinterpretation is open-ended rather than binary. To do so, we introduce the *MindLibs* task paradigm, in which we combine fMRI, eye-tracking, and behavioral measures while holding the physical stimulus constant before and after exposure to an alternative interpretation. Participants first generated their own interpretations of photographs depicting complex social scenes. They were then presented with alternative interpretations generated by other participants, ranging from highly semantically similar to highly distinct from their initial interpretation, and rated the extent to which they agreed with the initial versus alternative interpretation. This design allowed us to test whether reinterpretation was associated with conceptual updating, visual updating, or a coupling between the two. One possibility is that reinterpretation depends primarily on conceptual updating, such that people adopt a new interpretation by representing the stimulus differently in meaning-related neural systems such as the default mode network (DMN), which represents information in ways that depend on context, goals, and prior beliefs ^22–24^, even if visual sampling changes little. A second possibility is that reinterpretation depends on visual resampling: after encountering an alternative interpretation, participants direct their gaze differently, fixating on features that make the alternative construal more salient or compelling. Recent work shows that gaze behavior is tightly coupled to neural activity beyond visual cortex into systems associated with memory and higher-order cognition, including the DMN ^25–27^, raising a third possibility: that visual and conceptual updating may operate together, such that sampling the image differently helps support or stabilize an emerging shift in meaning.

Using a whole-brain, network-level approach, we find that reinterpretation is primarily associated with conceptual updating in lateral DMN regions, whereas visual resampling and associated changes in sensory regions contribute only insofar as they accompany this higher-order representational change.

## Results

### Task structure enables controlled and individualized exposure to alternative interpretations

To investigate how people update their interpretations of ambiguous information in response to an alternate perspective, we developed a novel task paradigm (*MindLibs*) that uses an adaptive experimental design. On each trial (**Fig. 1A**), participants (N = 56) viewed an ambiguous image (“initial viewing”, trial phase 1), generated an initial interpretation by filling in blanks of a predetermined scaffold (SELF; phase 2, see Methods, *Fill-in-the-blanks scaffold*), and rated their confidence in their interpretation (*Self_1_;* phase 3). On experimental trials (n = 30 trials), participants were then presented with an alternative interpretation provided by another participant (OTHER); on control trials (n = 15), they simply waited across a matched delay (phase 4). Participants then viewed the image again ("second viewing", phase 5) and rated their agreement with each interpretation (experimental trials, *Self_2_* and *Other*) or re-rated their own interpretation only (control trials, *Self_2_*; phase 6).

**Fig. 1 -.**
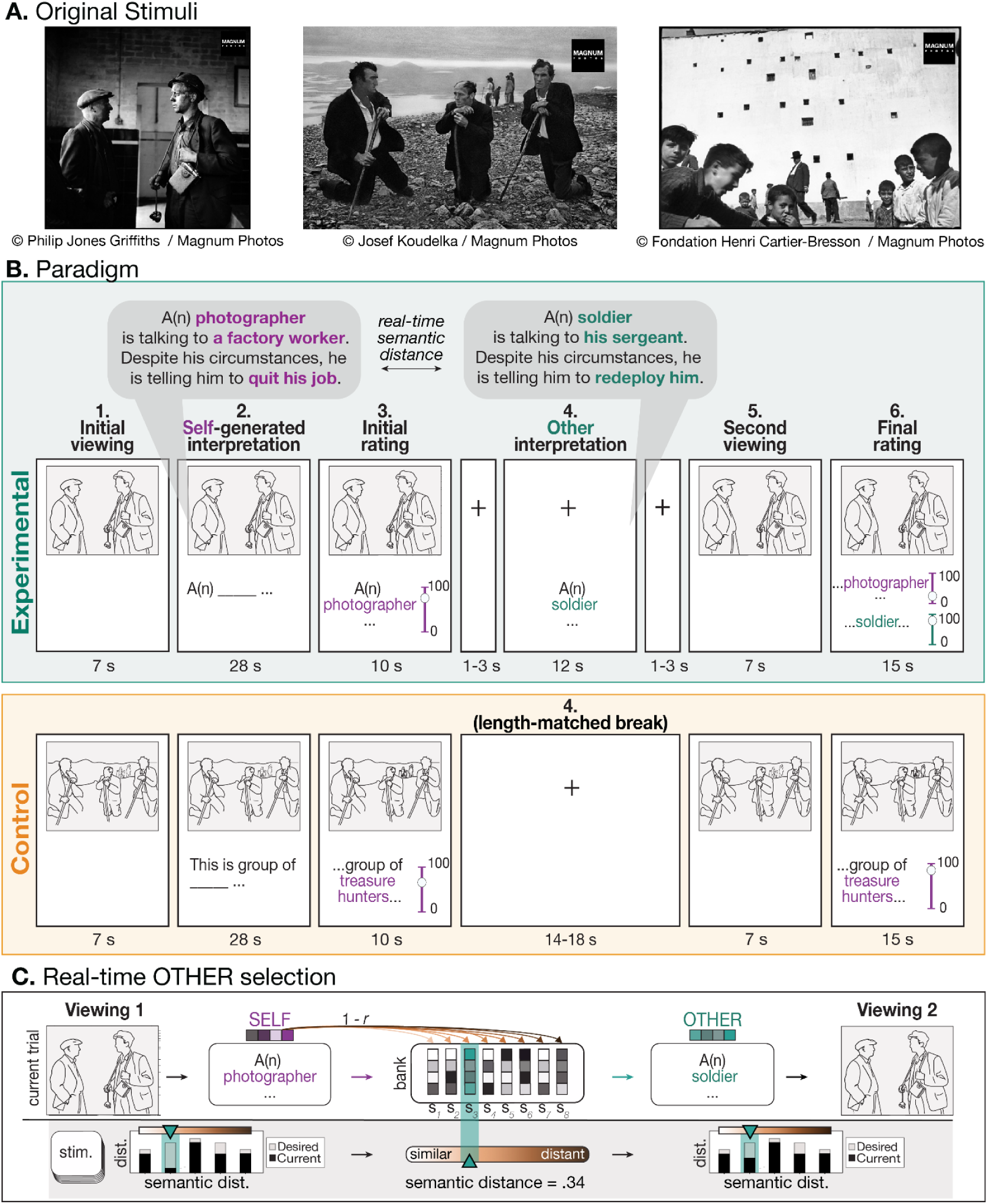
Paradigm. **A)** Sample original photographs used as experimental stimuli, reproduced here uncropped and with photographer and copyright attribution beneath each image. During the experiment, participants viewed versions without watermarks, cropped to a standardized size. **B)** Experimental paradigm. For illustrative purposes, the cropped photographic stimuli presented to participants are represented here as schematic line drawings created by the author (CSS); participants viewed the actual photographs, not the line drawings. Additional information on the stimuli is provided in **Supplementary** Fig. 1. In experimental trials, the alternative interpretations shown in Phase 4 were selected in real-time based on what the participant said during Phase 2 (see **A**). This was implemented by embedding the SELF using MiniLM-6^28^ and computing its semantic distance to a bank of eight candidate interpretations for each stimulus, sourced from previous participants. On each trial, the selected interpretation was drawn from the least-used quintile of the semantic distance distribution up to that point in the experiment and, where possible, from one of the least-sampled participants. Both semantic distance usage and participant sampling counts were updated dynamically across trials. See Methods, *Real-time “OTHER” selection,* for further details.

#### Image stimuli evoke variable interpretations

Each trial centered on a single ambiguous photograph (see **Fig. 1A** for an example; **Supplementary Figure 1** for additional examples). Ambiguity stemmed from uncertainty about the setting, interpersonal relationships, ongoing actions, and the emotions or mental states of the individuals depicted. Although each image presumably had a ground-truth context known by the photographer, this information was unknown to both experimenters and participants. For each image, we constructed a corresponding fill-in-the-blank scaffold, inspired by the classic children’s game *Mad Libs*, with three blanks that could be completed in different ways to produce alternative interpretations of the same image (see Methods, *Fill-in-the-blanks scaffold* for further details of stimulus selection and scaffold construction).

#### Trialwise real-time selection of OTHER based on semantic distance from SELF

On experimental trials, participants were presented with an alternative interpretation (OTHER) drawn from prior participants’ responses generated using the same scaffold. The semantic distance between SELF and OTHER, quantified as the distance between them in a high-dimensional semantic embedding space (miniLM-L6-v2 ^28^; see Methods, *Computing semantic distance*), served as the key manipulation. On each trial, OTHER was selected in real time to ensure that, across trials, each participant encountered alternatives ranging from highly similar to their own interpretation (distance ≈ 0) to highly divergent (distance ≈ 1; see **Fig. 1B**; Methods, *Real-time “OTHER” selection*). Alternatives were sourced from a pool of the eight most recently scanned participants; participants were told only that the alternative interpretation came from a previous participant (not necessarily the same individual across trials), with no further identifying information provided. This ensured sufficient coverage of the range of possible semantic distances to test the effects of semantic distance on neural, behavioral, and gaze shifts. Unless otherwise noted, all linear mixed-effects models reported below include random intercepts for participant and stimulus (image).

### Exposure to an alternative interpretation induces behavioral and neural but not gaze shifts

We first tested whether exposure to an alternative interpretation (OTHER)—independent of its semantic distance from the participant’s initial interpretation (SELF) or any resulting reinterpretation—elicited measurable changes in behavior, neural activity, and/or gaze.

#### Behavioral shifts

Participants changed their endorsement of their initial interpretation after being exposed to an alternative. Initial ratings (confidence; *Self_1_*; Task paradigm *Phase 3,* **Fig. 1**) did not differ between experimental and control conditions (β = 0.0001, *p* = 0.80, **Supplementary Fig. 2**), as expected since participants did not know which trial type they were in at this point. However, the absolute change from *Self_1_*to the subsequent self-rating (*Self_2_*; Task paradigm *Phase 6,* **Fig. 1**) was significantly larger on experimental than control trials (β = 0.025, p < 0.001; linear mixed effects model (LMEM) predicting |*Self_1_*– *Self*_2_| from the condition [experimental > control]; **Fig. 2A**).

**Fig. 2 -.**
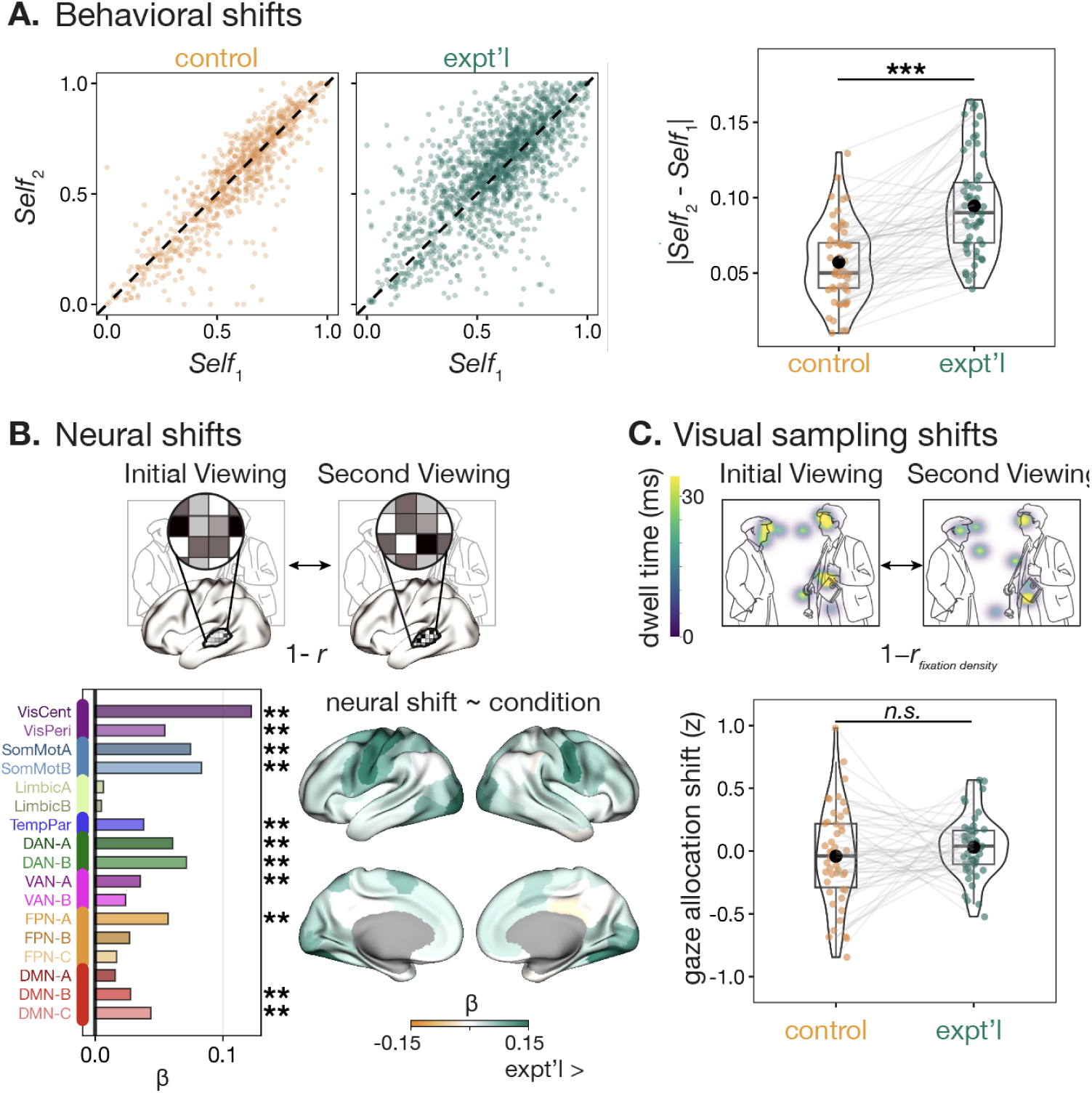
Neural and behavioral shifts occur in response to an alternative interpretation. **A)** *Left.* Trial-level Self₂ vs. Self₁ (confidence) ratings by condition. Each dot is one trial from one subject (n = 821 control trials; 1643 experimental trials). A greater change in self-endorsement (i.e., increased scatter both above and below the identity [dashed] line) in experimental trials can be appreciated visually. *Right.* Absolute difference between Self₁ and Self₂ for each participant in the experimental versus control conditions. Each dot represents the median across trials for a given participant (across 15 trials for the control condition and 30 trials for the experimental condition). Each gray line connects a participant to themselves. Experimental trials showed significantly more change in perceptions of one’s own interpretation than control trials (*p* < 0.001; LMEM). **B)** *Top.* Schematic of how neural shifts are computed. *Bottom.* Plotted estimates show the difference in neural shift magnitude between experimental and control trials. Estimates were obtained from LMEMs predicting within-subject neural shifts from condition (coded as experimental > control), with participant and stimulus included as random effects; network-level models additionally included parcel as a random effect. Analyses were matched for the number of trials in each condition (by subsampling 15 of the 30 total experimental trials); the median estimate from 100 iterations is plotted (see Methods, *Predicting neural shifts, Condition differences*). **C)** *Top.* Example gaze allocation during the initial and second viewing of a single trial, illustrating changes in visual sampling across viewings (see Methods, *Computing gaze allocation shifts*). Gaze data are from an actual trial, overlaid here on an author-created schematic representation of the original photographic stimulus for copyright purposes (see Fig. 1A). *Bottom.* Eyetracking results comparing gaze allocation shifts between conditions. Each dot represents the median across trials for a given participant. No significant differences were observed in how participants changed their sampling of the image between the initial and second viewing (*p* > 0.05; LMEM). Asterisks indicate significance (* *p* < 0.05, ** *p* < 0.01, *** *p* < 0.001; *q* in place of *p* for network-level analyses in B).

Thus, encountering an alternative interpretation reliably shifted participants’ endorsement of their original interpretation. These shifts could occur in either direction, as reflected by increases or decreases relative to initial ratings across trials (**Fig. 2A**). In experimental trials, on average, endorsement increased from the initial to the second rating (*Self*_2_ - *Self*_1_; estimated mean change = 0.031, SE = 0.006, 95% CI [0.018, 0.044], *p* < 0.0001), while in control trials, it did not change (estimated mean change = −0.011, SE = 0.007, 95% CI [−0.025, 0.004], *p* = 0.149). Exposure to an alternative interpretation thus reliably shifted endorsement, with updating biased toward strengthening rather than revising the original interpretation.

#### Neural shifts

We next examined whether neural activity showed analogous within-subject shifts following exposure to an alternative interpretation. For each trial, we estimated multivariate activity patterns (“neural representations”) in each of 100 cortical regions ^29^ during the Initial Viewing and Second Viewing (**Fig. 1A**, trial phases 1 and 5) — when, critically, visual stimulation was identical — and computed Fisher *z*-transformed Pearson correlations between them. Similarity of neural representations was highest in unimodal sensory and attention networks and progressively lower in transmodal association networks, including temporoparietal, default mode, and limbic systems (**Supplementary Fig. 3B**). To quantify changes in neural representations, we converted these correlations to distance values (“neural shifts”; see Methods, *Computing neural shifts*), where larger values indicate greater shifts between viewings. We then grouped parcels according to the 17-network cortical parcellation ^30^; **Supplementary Fig. 3A**), enabling analysis at the level of large-scale functional systems (network-level LMEMs included random effects of parcel).

Using these neural shift values, we compared experimental and control trials. Consistent with the behavioral findings (**Fig. 2A**), neural shifts were greater in experimental trials — i.e., following exposure to an alternative interpretation (whole-brain, *β* = 0.02, *p* < 0.001). All 17 Yeo networks showed effects in the predicted direction. While effects were strongest in both visual and somatosensory regions, they were also significant in all higher-order networks except DMN-A, limbic networks, and Frontoparietal control network (FPN)-C. At a finer spatial scale, most cortical regions exhibited this directional result, with the exception of the right posterior cingulate cortex, which showed no significant condition difference (**Fig. 2B**). This effect was robust to the trial number imbalance between conditions (30 experimental, 15 control), surviving correction in >99% of resampled analyses matching trial counts across conditions (see Methods, *Predicting neural shifts, Condition differences*). This suggests that changes in neural representation are in fact induced by exposure to an alternate interpretation, and are not due to simple re-exposure or adaptation effects.

#### Visual sampling shifts

Finally, we tested whether exposure to an alternative interpretation shifted how participants visually sampled the images using eyetracking data collected concurrently in the scanner. Gaze changes can be quantified in multiple ways; we began with a static measure of where gaze was allocated. We correlated duration-weighted fixation maps from the two viewings and converted this similarity measure to a distance (“gaze allocation shift”). This metric captures the extent to which gaze was concentrated in different regions of the image across viewings, irrespective of the order in which those regions were visited (see Methods, *Computing gaze allocation shifts*). We treated this as our primary measure of visual resampling.

However, because participants viewed the same image on both occasions, and because most images consisted of a small number of “attractors” (e.g., faces, salient objects), we reasoned that reinterpretation-related changes in gaze might be expressed not in how much time people spend looking at each spatial location, but in how they move between those locations over time. To capture this sequential structure, we computed scanpath dissimilarity using the “shape” metric from the MultiMatch algorithm, which quantifies differences in the geometric structure of gaze trajectories (^31,32^; see Methods, *Computing scanpath shifts*). Results using this measure are reported in the Supplement. The two measures were only modestly correlated (*r* = 0.33), suggesting that they capture largely distinct aspects of changes in viewing behavior.

Analyses were restricted to participants with usable eye-tracking data (*N* = 47). Although both gaze allocation shifts and scanpath shifts were numerically greater on experimental than control trials, neither difference was significant (gaze allocation shift: β = 0.005, *p* = 0.87; **Fig. 2C**; scanpath shift: β = 0.03, *p* = 0.36; Supplementary **Fig. 2C**).

Together, these findings show that exposure to an alternative interpretation reliably changes both behavioral endorsement of one’s original interpretation and neural representations of the stimulus, even before accounting for how distinct the alternative is, or whether it is ultimately adopted. Visual sampling, on the other hand, was not reliably altered by mere exposure to an alternative interpretation.

### When the alternative is highly distinct, neural but not gaze shifts reflect behavioral reinterpretation

Having established that exposure to an alternative interpretation changes behavior and neural activity, but not gaze, we next asked what determines the magnitude of these changes and whether they predict reinterpretation.

#### Participants are more likely to reinterpret when initial confidence is low and semantic distance is high

We first characterized the behavioral conditions under which participants were more likely to endorse the alternative over their own interpretation. Although participants tended to favor their initial interpretation (**Fig. 2A**, **Supplementary Fig. 2**), they sometimes endorsed the alternative. We quantified relative endorsement as the difference between final ratings of the alternative and self-generated interpretation (*Other* - *Self*_2_; **Fig. 1A**, Phase 6). Relative endorsement of OTHER depended on the interaction between initial confidence and semantic distance: participants were more likely to endorse an alternative when initial confidence (*Self*_1_) was low and semantic distance was high (β = -7.39, *p* < 0.001; **Fig. 3A**). (Note that semantic distance was modeled as a third-order polynomial to capture potential non-monotonic effects: here and throughout, the reported coefficient reflects the linear polynomial component and should not be interpreted as the effect of a one-unit change in semantic distance.)

**Fig. 3.**
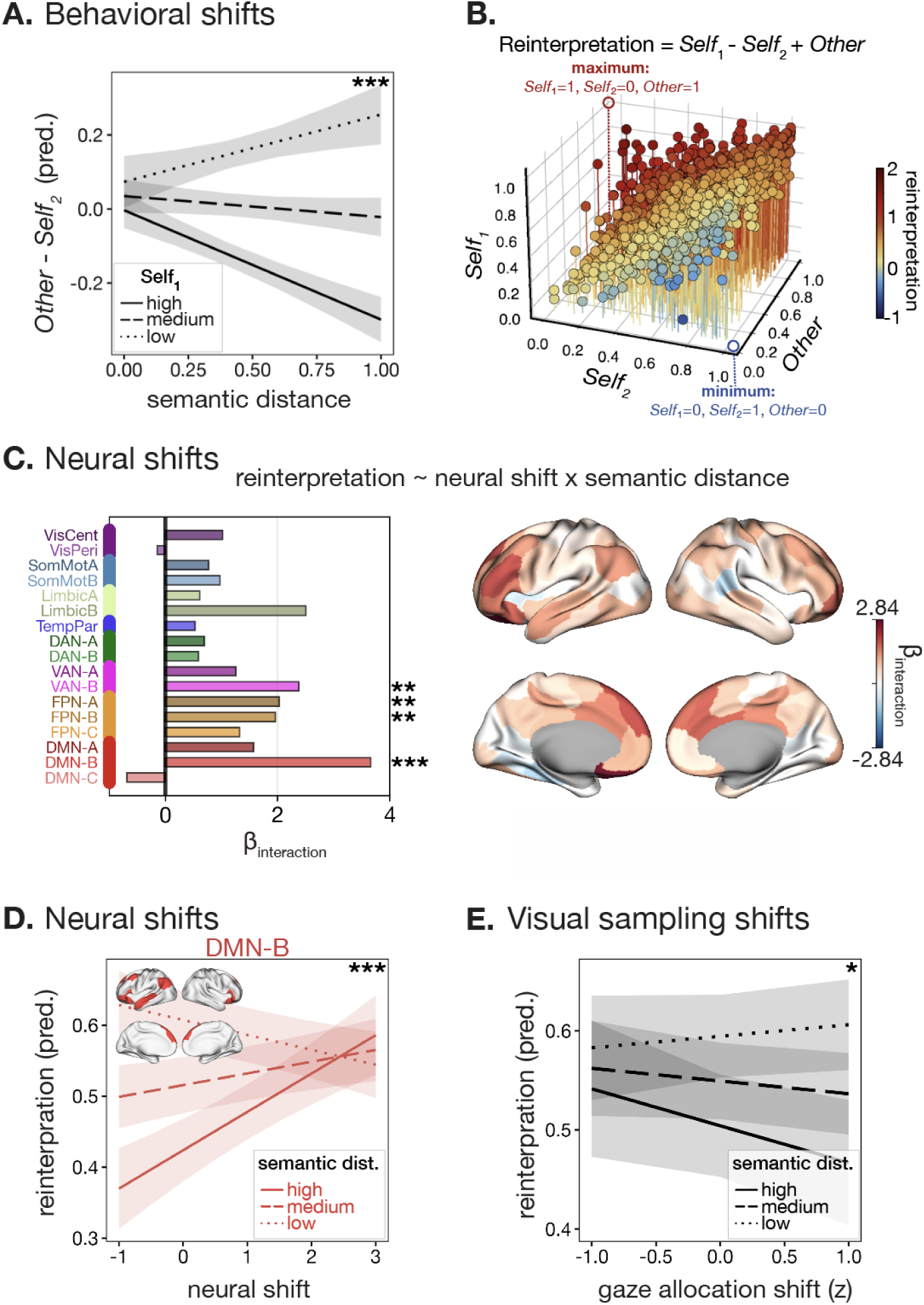
Participants hesitate to reinterpret, but when they do, neural shifts support that reinterpretation. **A)** Participants report more support for OTHER than for SELF at lower *Self*_1_ (index of confidence) and higher semantic distance. Predicted effects are shown from a LMEM estimating OTHER - SELF_2_ from an interaction between confidence and semantic distance. Dotted, dashed, and solid lines correspond to predicted values at confidence (Self_1_) values of 0.25, 0.50, and 0.75, respectively. **B)** Distribution of slider responses across all trials, colored by reinterpretation magnitude. Reinterpretation was quantified as *SELF_1_ − SELF_2_ + OTHER*. **C)** Output from linear mixed-effects models estimating reinterpretation from the interaction between neural shifts and semantic distance. *Left:* Network-level model estimates (including an additional random effect of parcel). *Right:* Parcel-level model estimates mapped onto the cortical surface (all models include random effects of subject and image). **D)** Predicted fixed effects from a linear mixed-effects model estimating reinterpretation from the interaction between neural shifts in DMN-B and semantic distance. The neural shift–reinterpretation association reversed sign with increasing semantic distance: negative at low distance (β = −0.021) and positive at intermediate distance (β = 0.0165), and high distance (β = 0.054); slopes differed significantly between low and high distance (*q* < 0.001). **E)** Predicted effects from a linear mixed-effects model estimating reinterpretation from the interaction between gaze allocation shifts and semantic distance Asterisks indicate significance (* *p* < 0.05, ** *p* < 0.01, *** *p* < 0.001; *q* in place of *p* for network-level analyses in C and D).

We therefore operationalize behavioral reinterpretation henceforth as *Self_1_ − Self_2_ + Other*, capturing both decreased endorsement of the initial interpretation and increased endorsement of the alternative (**Fig. 3B**).

#### Neural shifts support reinterpretation at high semantic distance

The behavioral results identify high semantic distance as the condition under which reinterpretation is most likely. To determine where neural shifts predict reinterpretation, we modeled reinterpretation as a function of neural shift, semantic distance, and their interaction. A whole-brain analysis including data from all parcels revealed a significant interaction between neural shifts and semantic distance (β = 4.46, *p* < 0.001): changes in how the stimulus was neurally represented predicted behavioral reinterpretation selectively at high semantic distances. Effects were strongest in lateral regions of DMN-B (e.g., left temporoparietal junction), followed by FPN-A and B (e.g., dorsolateral PFC), and VAN-B (e.g., anterior cingulate; **Fig. 3C, D**). Results were robust across semantic embedding models (MiniLM-L12, MPNet ^28,33^) and alternative operationalizations of reinterpretation (**Supplementary Fig. 4A**).

Across networks showing significant interactions, the association between neural shifts and reinterpretation flipped sign with increasing semantic distance: negative at low distance, absent at intermediate distance, and positive at high distance. This was clearest in DMN-B, where slopes differed significantly between low and high distance (Δβ = −0.074, SE = 0.019, *p* < 0.001; **Fig. 3D**). This suggests that stable neural representations help maintain an initial interpretation when alternatives are semantically close, whereas greater representational change supports reinterpretation when alternatives are more distant.

#### DMN-B carries an independent reinterpretation-related signal

The network-level analyses identified several higher-order association network—DMN-B, FPN-A, FPN-B, and VAN-B—in which neural shifts predicted reinterpretation as a function of semantic distance, with DMN-B showing the strongest effect. We next asked whether these networks contribute unique variance to reinterpretation, or whether some are largely reducible to variance shared with DMN-B. To test this, we fit mutual adjustment models estimating each network’s neural shift x semantic distance interaction while controlling for the other networks (see Methods, *Assessing shared variance between DMN-B and other association networks*). DMN-B’s effect attenuated only modestly after controlling for the other networks, whereas effects in FPN-A, FPN-B, and VAN-B were each substantially reduced after controlling for DMN-B (**Supplementary Fig. 4B**). This asymmetry suggests that FPN and VAN effects are not independent of DMN-B; rather, DMN-B appears to carry the more robust, dissociable signal linking neural updating to behavioral reinterpretation.

#### Visual sampling shifts do not predict reinterpretation

We next tested whether shifts in visual sampling predicted reinterpretation. There was no main effect for either measure (gaze allocation, *p* = 0.62; scanpath, *p* = 0.67). Unlike neural shifts (**Fig. 3D**), neither measure showed the positive interaction with semantic distance that would be expected if visual resampling drove reinterpretation. For gaze allocation, the interaction was statistically significant but in the opposite direction (β = −0.59, *p* = .049; **Fig. 3E**), and did not survive correction for the two measures tested; for scanpath, it was absent (β = 0.14, *p* = 0.64; **Supplementary Fig. 6C**). This pattern is consistent with the preceding condition analysis, in which visual sampling shifts did not reliably differ between experimental and control trials. Together, these findings suggest that, unlike conceptual updating, visual resampling alone is not strongly related to reinterpretation.

Altogether, these results show that it is precisely when alternatives are most distinct that neural shifts become predictive of reinterpretation, and that this relationship is largely specific to conceptual rather than visual resampling: DMN-B’s representational changes track the active adoption of a new interpretation, while shifts in visual sampling showed little relationship to reinterpretation on their own. If gaze alone was not sufficient to predict reinterpretation, might it still matter in combination with conceptual updating in DMN-B? We next tested if and how the coupling between conceptual and visual resampling relates to reinterpretation.

### Visual resampling does not drive reinterpretation-linked neural shifts in DMN-B

While gaze shifts alone were not sufficient to predict reinterpretation, it is still possible that visual resampling plays a role in combination with conceptual updates; we investigate this possibility in our final set of analyses.

#### Visual sampling shifts do not interact with the DMN-B reinterpretation signal

We first tested whether the magnitude of visual sampling shifts moderated the relationship between DMN-B neural shifts and reinterpretation, as a function of semantic distance. We extended our model to include the three-way interaction among neural shift, gaze allocation shift, and semantic distance (see Methods, *Predicting reinterpretation*). The neural shift x semantic distance interaction remained significant (β = 3.31, *p* < 0.001), but the three-way interaction with gaze allocation shift, while trending in the expected direction, did not reach significance (linear β = 1.36, *p* = 0.052). At high semantic distance, the positive relationship between DMN-B neural shifts and reinterpretation was present across the full range of gaze allocation shifts, from low (β = 0.078, *p* <0.001) to moderate (β = 0.089, *p* < 0.001) to high (β = 0.10, *p* < 0.001; **Fig. 4A**), with no significant difference between low and high (Δβ = 0.022, p =0.49). Analyses using the dynamic scanpath measure to quantify visual shifts showed similar results (**Supplementary Fig. 5**). Thus, the relationship between DMN-B neural shifts and reinterpretation did not reliably vary with visual resampling.

**Fig. 4.**
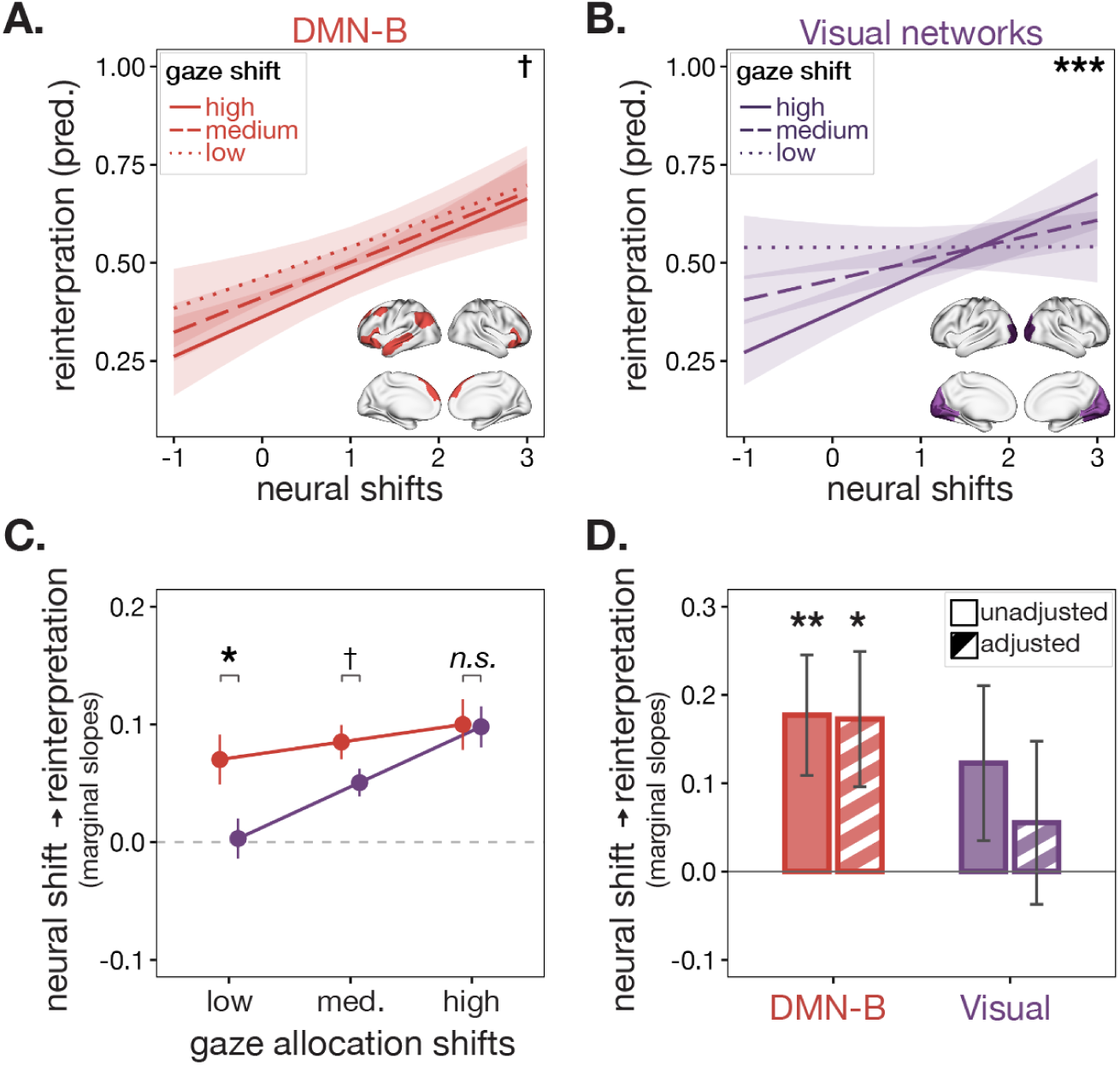
DMN-B neural shifts predict reinterpretation independently of visual resampling, whereas visual-network effects depend on gaze. Predicted fixed effects from a linear mixed-effects model estimating reinterpretation from the interaction between neural shifts, gaze allocation shifts, and semantic distance (random effects of subject and image; see Methods, *Predicting reinterpretation*). Effects are plotted at high semantic distance (0.75). Low, medium, and high gaze allocation shifts correspond to −1, 0, and +1 SD from the mean (z-scored within participant). Statistics reflect comparison of slopes for the interaction between neural shifts and gaze allocation shifts at high semantic distance. **A)** Predicted fixed effects in the DMN-B network, highlighting that neural shifts predict reinterpretation at high semantic distance independently of gaze. **B)** Predicted fixed effects in the visual networks, highlighting that neural shifts predict reinterpretation at high semantic distance specifically when gaze allocation shifts are large, consistent with a sensory role contingent on active visual resampling. **C)** Estimated marginal slope of neural shifts on reinterpretation (at high semantic distance) across low, medium, and high gaze allocation shifts (as in A and B), for DMN-B and the visual networks. DMN-B’s slope is steeper than visual at low and moderate gaze shifts, but not at high gaze shifts. Error bars reflect ±1 SE. **D)** Marginal slope relating each network’s mean neural shift to reinterpretation at high semantic distance and high gaze shift, before (solid) and after (hatched) adjusting for the other network (see Methods, *Assessing Shared Variance Between DMN-B and visual networks*). Error bars reflect ±1 SE. This further indicates that DMN-B’s relationship with reinterpretation is robust and independent, whereas visual effects are attenuated after controlling for DMN-B. Asterisks indicate significance (* *p* < 0.05, ** *p* < 0.01, *** *p* < 0.001, , ✝ *p* < 0.10, *n.s.* = not significant).

#### Visual-network effects depend on gaze and are not independent of DMN-B

To evaluate whether this pattern was specific to a network implicated in conceptual processing, we repeated the analysis in visual cortex. The central visual network (VisCent), comprising primary visual regions around the calcarine sulcus, showed the largest effect size in the condition analysis comparing neural shifts between experimental and control trials (**Fig. 2B**); we reasoned that any reinterpretation-related effects attributable to visual resampling should be especially apparent in this network. For simplicity, we combined the central (VisCent) and peripheral (VisPeri) visual networks into a single visual-network measure, given their common assignment to the visual system in the Schaefer/Yeo 17-network parcellation. Effects for each network separately are shown in **Supplementary Fig. 6**.

Neural shifts in either visual network did not predict reinterpretation in the two-way model (**Fig. 3**), but became related to reinterpretation once gaze allocation shift was included (three-way interaction; linear β = 2.18, *p* = 0.001). Unlike DMN-B, this positive relationship was significantly greater at high levels of gaze allocation shift (Δβ comparing high to low = 0.10, *p* < 0.001), indicating that for visual networks, visual resampling was necessary for neural shifts to predict reinterpretation (**Fig. 4B**). To formally test this dissociation, we compared the two networks’ estimated marginal slopes directly: DMN-B’s slope was steeper than the visual networks’ at low gaze allocation shift (Δβ = 0.07, *p* < 0.05), did not differ reliably at moderate shift (Δβ = 0.03, *p* = 0.06), and was indistinguishable at high shift (Δβ = 0.001, *p* = 0.94; **Fig. 4C**; see Methods, *Comparing DMN-B and visual networks*). Thus, reinterpretation-related shifts in visual networks emerged only when participants substantially changed how they sampled the image.

Finally, we asked whether the reinterpretation-related signals in DMN-B and the visual networks were independent, or whether one was largely reducible to variance shared with the other (see Methods, *Assessing shared variance between DMN-B and visual networks*). For each network we estimated the slope relating its neural shift to reinterpretation at high semantic distance, before and after entering the other network into the same model. As predicted, DMN-B’s slope was essentially unchanged by adjustment for the visual networks (slope = 0.177, unadjusted; slope = 0.175, adjusted; 1.3% attenuation, both *p* < 0.05). In contrast, the visual network’s slope was largely attenuated (slope = 0.119, *p* = 0.173, unadjusted; β = 0.050, *p =* 0.585, adjusted, 57.8% attenuation; **Fig. 4D**).

This pattern was consistent in both independent visual networks and was stronger in VisPeri than VisCent (**Supplementary Fig. 6**). Further, all of these results generalized to using scanpath shifts as the visual resampling metric (**Supplementary Fig. 5**).

Together, this pattern of results suggests that visual resampling and associated shifts in visual cortex accompany reinterpretation but do not independently account for it. Instead, reinterpretation appears to be associated primarily with higher-order conceptual updating in DMN-B, with visual changes playing a conditional, secondary role.

## Discussion

The present study investigated how individuals revise their interpretations of ambiguous stimuli in response to alternative meanings. Using a novel paradigm that combined self-generated interpretations with dynamically selected alternatives, we found that conceptual and visual updating showed dissociable relationships with reinterpretation. Shifts in neural representations of the stimulus in lateral default mode regions (DMN-B) predicted reinterpretation, particularly when the alternative interpretation was semantically distant, independently of neighboring frontoparietal and attention networks and regardless of changes in gaze. In contrast, visual resampling did not independently predict reinterpretation. Instead, neural shifts in visual networks tracked reinterpretation only when accompanied by substantial gaze shifts. Critically, this gaze-gated visual contribution did not account for unique variance beyond shifts in DMN-B, whereas the DMN-B effect was unchanged when visual and gaze terms were included. Together, these results suggest that the primary route to reinterpretation is conceptual updating linked to representational change in higher-order regions, with visual resampling playing a conditional and largely redundant role.

These findings build on evidence that the way we experience information is not a fixed readout of sensory input but is shaped by memory, affect, motivation and self-relevant meaning ^34–36^. Prior paradigms investigating how ambiguous stimuli are interpreted and reinterpreted have varied along several dimensions, including whether interpretations are fixed versus open-ended, self-generated or externally imposed, and whether the physical stimulus is held constant or dynamically unfolding. Studies using perceptual disambiguation (e.g., Mooney images) typically hold the stimulus constant and show that priors that impose a single, fixed interpretation sharpen and reshape neural representations within frontoparietal and/or default mode networks ^37,38^. While visual sampling may change following disambiguation ^39^, its contribution when perceptual ambiguity persists — e.g., in the case of bistable images with two equally plausible percepts — is less clear ^40^. Studies using dynamically unfolding narrative stimuli show that default mode representations differ across individuals who interpret the same information differently ^23,41–43^, and change within an individual when new information is introduced that changes the dominant interpretation ^22,24^. Given that sensory input unfolds along with the interpretation, however, these paradigms provide less leverage for determining whether changes in sensory sampling drive and/or accompany reinterpretation. *MindLibs* builds on both of these traditions: we hold the stimulus constant, making it straightforward to dissociate changes in interpretation from changes in the stimulus, while having participants compare between two theoretically plausible alternatives that differ in their source (self versus someone else). This allows us to track both representational change across the brain and visual resampling as participants decide whether to move from a self-generated interpretation toward a competing alternative. Critically, the magnitude of representational change in association cortex, but not visual resampling, scaled with the degree to which participants reported adopting the alternative. Representational change in visual cortex predicted reinterpretation only when accompanied by large gaze shifts, suggesting that shifts in sensory regions reflect changes in visual input produced by changes in visual sampling rather than changes in the subjective percept of that input *per se.* This distinction is consistent with accounts that separate changes in perceptual input from genuine top-down effects of cognition on perception ^44^.

Prior approaches also differ in whether reinterpretation serves an instrumental role or is treated as an outcome in its own right. Cognitive reappraisal paradigms, for example, instruct participants to adopt a new construal in service of an explicit affective goal and still implicate the a similar set of regions (e.g., ^45,46^). That these functionally distinct forms of reinterpretation converge on an overlapping spatial pattern raises the possibility of a common mechanism for revising the meaning of a stimulus when multiple interpretations remain viable. Our findings may also help contextualize mixed evidence linking gaze behavior to reappraisal success ^47–50^ by showing that gaze changes are not uniformly necessary for updating the meaning of a stimulus.

This paradigm further offers two methodological advances beyond the dimensions above. First, by using participants’ own open-ended interpretations and similarity scores derived from language models to adapt trials in real time, we define each participant’s interpretive space using their own responses, rather than a fixed set of options or a normative baseline. This allows interpretive distance to be manipulated within individuals, trial by trial, without assuming what the "correct" interpretation should be. This approach could extend to studying belief updating, social reasoning, and conceptual change more broadly, in a way that is both controlled and sensitive to individual differences. Second, holding the physical stimulus constant across viewings allowed us to more cleanly attribute neural and gaze shifts to changes in a participant’s own construal of the image, rather than to changes in the input itself. Since the stimulus never changed, any shift in visual sampling reflects a change in how participants explored the image in light of a new interpretation, rather than a response to new visual content. Gaze and neural activity are tightly coupled during naturalistic viewing ^26^, such that a continuously unfolding stimulus can drive both simultaneously, making it difficult in dynamic paradigms to isolate conceptual from visual updating. By holding the image fixed, our design removes stimulus-driven change and allows us to more directly dissociate these two possible routes to reinterpretation.

Still, several limitations should be considered. First, while distance in the semantic embedding space of a large language model provides a useful operationalization of interpretive variability, it may not capture other relevant dimensions, such as emotional salience or social relevance. Future work could manipulate these dimensions directly, for example by varying the valence ^3^ of alternative interpretations or the identity and reliability of their source. Second, the present analyses focus on regional multivariate patterns; future work could take more time-resolved approaches to attempt to dissociate the neural dynamics underlying the decision to adopt an alternative interpretation from its implementation. Relatedly, although the present analyses distinguish conceptual representational shifts from visual resampling, they do not establish their temporal ordering. Conceptual updating in DMN-B could precede visual resampling, follow it, or interact recurrently with visual regions through top-down modulation ^13^. Similar questions of directionality remains unresolved even in related domains: studies of gaze–neural coupling during memory retrieval report similar ambiguity, with evidence for both hippocampal activity driving gaze ^51^ and the reverse (^52^; see ^53^ for a recent review). Likewise, because gaze indexes only overt sampling, our measures cannot detect a change in meaning accompanied by covert attentional reallocation. Since the image was held constant across viewings, however, reinterpretation in the absence of gaze changes still reflects a change in meaning that cannot be attributed to changes in overt visual sampling, even if covert attention reallocation contributed to that change.

In conclusion, reinterpretation under persistent ambiguity is a structured and constrained process, shaped by semantic distance between the original and alternative interpretations and expressed as a measurable reorganization of the stimulus’s neural representation. By preserving ambiguity and allowing participants to decide whether to adopt an alternative interpretation, we show that reinterpretation is carried primarily by changes in higher-order conceptual representations in DMN-B, largely independent of how participants sampled the image. Gaze itself did not drive reinterpretation; rather, gaze shifts set the conditions under which a relationship with visual-cortex representations emerged, suggesting that while reinterpretation is at its core conceptual, it can be optionally accompanied by visual sampling changes. More broadly, these findings suggest that when no single interpretation is definitively correct, revising meaning depends less on how we look than on how we represent what we see.

## Methods

Participants completed a novel paradigm — the "MindLibs" task — in which they generated open-ended interpretations of ambiguous photographs and then evaluated an alternative interpretation provided by a previous participant. Participants completed this task in the MRI scanner with simultaneous eye-tracking (details of data acquisition below). We quantified the semantic distance between interpretations using sentence-level embeddings, neural responses using multivariate pattern analysis, and eye movements using gaze metrics. We then examined how these signals (individually and in combination) predicted the degree to which participants updated their interpretations.

### Participants

The Committee for the Protection of Human Subjects of Dartmouth College approved all data collection, and all participants provided written informed consent prior to participation.

### fMRI participants

Participants (n = 56; 23 male; median age = 20, range = 18-39) were healthy adults recruited from the local areas of New Hampshire and Vermont, including the Dartmouth College student body. All participants had normal or corrected-to-normal vision and hearing and reported no psychiatric or neurological diagnoses or MRI contraindications. Of 62 initially recruited participants, 3 were excluded due to equipment malfunction and 3 due to excessive head motion (framewise displacement > 0.2 mm), yielding the final sample of 56. Within the remaining participants, head motion was low (mean = 0.099, SD = 0.108). To ensure data consistency, trials with excessive motion were excluded from both imaging and behavioral analyses (see f*MRI data acquisition and processing*).

### Stimulus Set

All images were sourced from Magnum Photos (magnumphotos.com) in November 2021. Each photograph was manually cropped—while preserving all primary subjects and narrative features—to ensure a consistent display size of 2.5 x 2.0 inches (≈ 6.35 x 5.08 cm). The final stimulus set consisted of 45 black-and-white documentary photographs drawn from nine major Magnum photographers: Henri Cartier-Bresson (n=6), Werner Bischof (n=5), Stuart Franklin (n=4), Josef Koudelka (n=4), Elliott Erwitt (n=6), Philip Jones Griffiths (n=9), Carl De Keyzer (n=6), Constantine Manos (n=2), and David Hurn (n=3). These images were selected from a larger candidate pool on the basis of their ability to evoke a broad range of open-ended interpretations in pilot testing.

All stimuli depict ambiguous social scenes lacking a canonical or normative interpretation. Across images, participants typically diverged in how they construed (i) the setting, (ii) the relationships among individuals, (iii) the implied actions or events, and/or (iv) the mental states or emotions of the characters.

### Fill-in-the-blanks scaffold

Each image was paired with a custom “Mad Libs”-style fill-in-the-blanks prompt. These prompts were designed to capture meaningful interpretive variability across participants while holding fixed the more stable, consensus elements of each photograph (e.g., “a group” versus “a group of friends/people/enemies”). Prompts were created in a three-step process. (1) Forty participants completed a survey in which they freely described what they observed in each image. Instructions read: “*Please describe what is happening in this image in 2–4 sentences. Focus on who is in the image and their thoughts and actions. Write at least 20 words for each response*.” A description for a sample image was provided as guidance. (2) These narratives were analyzed to identify (i) areas of consensus (e.g., the number of people present or the general setting) and (ii) areas of ambiguity where interpretations diverged most strongly. Based on this analysis, we constructed a unique three-blank prompt for each image, with blanks targeting precisely those ambiguous elements—typically the people’s identities, locations, emotions, intentions, or relationships. The scaffolds were drafted manually by C.S.-S. and reviewed by E.S.F. Although every prompt contained three blanks, the total prompt length varied across images (M = 18.2, SD = 3.9 words, range = 12–30). Blanks did not specify the required part of speech. (3) We then tested these prompts with an expanded sample of participants (N > 500) and computed pairwise semantic similarity between all responses generated for each image. The 45 images that elicited the greatest interpretive variability—indexed by the lowest median cosine similarity across pilot responses—were selected for inclusion in the fMRI study.

### "MindLibs" Task Paradigm

Participants completed two sessions, one week apart; this manuscript uses data from the first session only. Each trial consisted of the following sequence: (1) an initial image viewing (7 s); (2) a self-generated interpretation via the fill-in-the-blanks prompt ("SELF", 28 s), which participants spoke aloud and was typed into the task-presentation computer in real time by the experimenter; (3) an endorsement rating of the self-generated interpretation (*Self_1_*, "confidence"; 10 s); (4) presentation of an alternative interpretation ("OTHER"; experimental trials only) or viewing a fixation cross across a matched-length delay (control trials; 12 s); (5) a second image viewing (7 s), and finally (6) an endorsement rating for both the SELF and OTHER interpretations (*Self_2_* and *Other*, experimental trials) or just the SELF interpretation again (*Self_2_,* control trials, 15 s). Ratings were made on independent sliders ranging from 0% ("strongly disagree") to 100% ("strongly agree"). Participants were told that there were no correct answers, that the OTHER interpretations were sourced from previous participants (not necessarily the same person across trials), and that the sliders were independent and did not need to sum to any particular value. In the MRI, jittered fixation intervals (1–3 s) were inserted to optimize event-related design efficiency and improve deconvolution of hemodynamic responses (**Fig. 1A**).

Across the experiment, participants completed 30 experimental trials and an additional 15 control trials, with image–condition assignment randomized across participants. In experimental trials, participants were instructed to read the OTHER interpretation aloud to promote engagement and match encoding demands as best as possible between the SELF and OTHER interpretations. While speaking in the scanner can increase head motion, our primary periods of interest were trial phases 2 and 5, during which participants were silent; our preprocessing strategy was chosen to further mitigate any residual effects of head motion (see *fMRI data acquisition and processing* below for more details).

### Computing semantic distance

To quantify the semantic content of participants’ interpretations, we computed sentence-level embeddings that incorporated both the shared scaffold of each prompt and the participants’ fill-in-the-blank responses. Each scaffold prompt consisted of three fixed text segments (“mindlib” – ML1, ML2, ML3) separated by three blanks (inputs 1–3), and the semantic embedding for each blank was computed within its appropriate linguistic context. This ensured that embeddings reflected the meaning of a participant’s response *as used within the prompt*, rather than as an isolated word or phrase.

For the first blank, we embedded the participant’s response within the context of the opening scaffold (ML1). For the second blank, the embedding was computed using the accumulated context up to that point (ML1 + input1 + ML2), and similarly, the third blank was embedded within the full preceding context (ML1 + input1 + ML2 + input2 + ML3). For each blank, we identified the tokens corresponding specifically to the participant-generated text and extracted the corresponding contextualized embedding vectors from the MiniLM-L6 model ^28,54^. Multi-token responses were summed into a single vector to yield one embedding per blank. This method preserves the compositional structure of the prompts and accounts for the evolving linguistic context across blanks.

To compute semantic similarity between two interpretations, we averaged the three blank-specific embeddings to obtain a single composite embedding per interpretation. Pairwise semantic similarity (e.g., SELF vs. OTHER) was quantified using cosine similarity between composite embeddings; we use the term semantic distance to refer to the corresponding cosine distance (1 − cosine similarity).

### Real-time “OTHER” selection

To ensure that participants encountered OTHER interpretations spanning the full range of semantic similarity, we selected OTHER interpretations in real time during fMRI scanning. During each trial, the participant’s SELF interpretation as typed by the experimenter was immediately converted into a sentence embedding using the MiniLM-L6 model, chosen for its computational efficiency ^28^. Pairwise similarity values were binned into five quintiles: very similar (0.8–1.0), similar (0.6–0.8), moderate (0.4–0.6), dissimilar (0.2–0.4), and very dissimilar (0.0–0.2). The current participant’s embedding was compared against a rolling pool of interpretations from the eight most recently scanned participants, classified into quintiles, and an OTHER interpretation was selected from the quintile least represented in earlier trials, ensuring balanced sampling across the similarity spectrum (**Fig. 1B**). When possible, the algorithm also maintained relatively even representation across source participants. When no suitable OTHER interpretation could be identified within the target range, the system selected a random interpretation from a larger database of pilot responses to ensure an OTHER interpretation could always be presented without delay. Across the experiment, semantic distances spanned a broad range (M = 0.42, SD = 0.19, range = 0.00–0.90).

### Computing reinterpretation

Reinterpretation was operationalized as the degree to which participants shifted away from their initial interpretation and toward the alternative, combining both decreased endorsement of the self-generated interpretation and increased endorsement of the alternative. Formally, reinterpretation was computed as: *reinterpretation = Self_1_ − Self_2_ + Other,* where *Self_1_* is the initial confidence rating (**Fig. 1A**, phase 3), *Self_2_*is the final rating of the self-generated interpretation, and *Other* is the final rating of the alternative interpretation (**Fig. 1A**, Phase 6). Higher values thus reflect greater updating: a combination of reduced confidence in one’s initial interpretation and greater endorsement of the alternative.

To validate our operationalization, we performed a principal components analysis (PCA) on the three behavioral ratings (*Self*_1_, *Self*_2_, *Other*), after z-scoring within participant. The first two principal components explained 91% of the variance in the appraisal space. PC1 reflected overall appraisal magnitude, with positive loadings from all three measures (*Self*_2_ = 0.66, *Self*_1_ = 0.63, *Other* = 0.40). PC2 captured a perspective contrast, with a large positive loading for other-appraisal (*Other*= 0.91) and smaller positive loadings for self-appraisal and confidence (*Self*_2_ = 0.32, *Self*_1_ = 0.25). Our reinterpretation measure correlated more strongly with PC2 (*r* = 0.76) than PC1 (*r* = 0.36), confirming that *Self*_1_ − *Self*_2_ + *Other* captures a genuine dimension of perspective-shifting in the data rather than global response intensity.

We additionally considered alternative operationalizations of reinterpretation, including a measure emphasizing relative support for the alternative interpretation (*Other* / (Self_2_ + *Other*)), which showed a commensurate relationship with our neural data (**Supplementary Fig. 4A**).

### Eyetracking data preprocessing

Raw gaze data were recorded using an MR-compatible SR Research EyeLink 1000 Plus device. A 9-point calibration was performed before each functional run using a Pygame calibration display passed to the EyeLink host via `pylink.openGraphicsEx.` Data was collected in EyeLink Data Format (EDF) and converted to ASCII using the edf2asc command-line tool (SR Research Ltd). Data from each run were parsed with custom Python scripts that extracted gaze samples, fixation/saccade events, and task messages separately. Samples flagged as tracker loss (off-screen sentinel values returned during blinks or signal loss) were removed, and remaining samples were retained only if they fell within the screen boundaries (0–1920 x 0–1080 px).

During the inter-trial intervals, participants fixated a central cross, so gaze during these periods should be stable. We used these intervals to correct for slow between-run drift, shifting each run so that its median inter-trial gaze position matched the median across all runs. Corrected samples were then restricted to those falling within the boundaries of the stimulus image, extended by a 100-pixel margin to accommodate residual calibration offset; gaze beyond this margin was excluded.

Eyetracking data quality is known to be substantially degraded in the MRI environment due to head coil interference, restricted head movement, and scanner vibrations. Given that both of our measures are computed from discrete fixation events rather than raw gaze samples (see *Computing gaze allocation shifts*), we expected them to be relatively robust to high-frequency sample noise; nonetheless we applied trial-level quality control. Off-image gaze was removed at the sample level, and trials for which a comparison could not be computed were excluded automatically. The two measures differ slightly in their minimum requirements—the fixation density map (“gaze allocation shift”) requires at least one fixation per viewing, whereas MultiMatch (“scanpath shift”) requires at least two after simplification—so the number of usable trials differs modestly between them (1,280 and 1,228, respectively). We then did a manual inspection to target artifacts that these automatic filters could not capture. Specifically, an experimenter visually inspected each trial’s gaze trajectory and marked trials for exclusion on the basis of data quality (e.g., tracker-dropout streaks or a frozen, flat-lined signal with no gaze variation across a viewing). This inspection was performed blind to the behavioral measures. For the analyses reported here, the eyetracking data were restricted to the trials retained in the fMRI dataset (i.e., trials whose corresponding image passed fMRI motion QC), yielding a final sample of 47 participants (*M* = 24.89 trials per participant, range = 5–42). Data from the majority of participants (N = 32) was collected at 250 Hz; a subset (N = 15) was collected at 1000 Hz. All effects withstood subsetting for either group.

### Computing gaze allocation shifts

Fixation events during the two viewings were identified using the EyeLink online parser (SR Research); saccades and blinks were excluded prior to analysis. Consecutive fixation-classified samples separated by no more than 5 ms were grouped into discrete events, each defined by its mean location (x, y) and duration, and events comprising fewer than two samples were discarded. Fixations interrupted by a gap of ≤ 150 ms and separated by ≤ 50 px were then merged into a single duration-weighted event, recovering fixations fragmented by blinks or brief tracker dropout, and the resulting events shorter than 100 ms were discarded. The same fixation events were used for both measures reported below.

For each viewing, we constructed a duration-weighted fixation density map. The image region —the stimulus bounds extended by the 100-px margin described above, spanning 950 x 800 px —was divided into a 50 x 50 grid (cells of 19 x 16 px, or 0.50° x 0.42° of visual angle). Each fixation contributed its duration to the cell containing it, so that cells accumulate dwell time rather than fixation counts. The resulting map was smoothed with a Gaussian kernel (σ = 1 grid cell). Gaze allocation similarity was quantified as the Pearson correlation between the two viewings’ smoothed maps across all 2,500 cells. Gaze allocation shift was computed as 1 − *r*, so that higher values indicate a greater change in how gaze was distributed across the image. Values were z-scored within participants across all valid trials.

This measure captures whether gaze was concentrated in the same regions, and for the same duration, in both viewings. It is insensitive to the order in which regions were visited: two viewings sampling identical locations in a completely different sequence would yield a high correlation. This property motivated the complementary scanpath measure described below, which is sensitive to sequence but not to absolute location.

### Computing scanpath shifts

Scanpath shifts were computed from the same fixation events. Fixation sequences were passed to the MultiMatch package ^31,32^, which derives saccade vectors between consecutive fixations, simplifies each scanpath by merging successive saccades differing by less than 45° or shorter than 75 px, aligns the two simplified scanpaths via their vector differences, and quantifies their similarity along five dimensions—shape, direction, length, position, and duration—each on [0, 1] (1 = identical). We chose the shape metric from the MultiMatch toolbox, which indexes the overall geometric form of the scanpath based on the vector difference between corresponding saccades, normalized by the screen diagonal (cf. ^55^, who note that the choice of scanpath measure must be hypothesis-driven). We hypothesized that adopting an alternative construal of an unchanged scene might reorganize how participants visually traverse an image, producing a corresponding change in scanpath geometry. Shape offers a second advantage particular to the MRI environment. Since it is computed from saccade vectors rather than absolute fixation coordinates, it is unaffected by translation of the scanpath as a whole, in which every fixation shifts by the same offset while the vectors between them are preserved, as might occur with calibration drift or head displacement.

Scanpath shift was computed as 1 − shape similarity, so that higher values indicate greater reorganization of the movement pattern between viewings. Values were z-scored within participants across all valid trials.

### fMRI data acquisition and processing

#### MRI acquisition

All data were collected at Dartmouth College using a 3.0 Tesla Siemens MAGNETOM Prisma whole-body MRI system (Siemens Medical Solutions, Erlangen, Germany) equipped with a 64-channel head coil.

#### T1 image

For registration purposes, a high-resolution T1-weighted magnetization-prepared rapid acquisition gradient echo (MPRAGE) imaging sequence was acquired (TR = 2,300 ms, TE = 2.32 ms, inversion time = 933 ms, flip angle = 8°, matrix size = 256 x 256, slices = 192, voxel size = 0.90 x 0.94 x 0.94 mm). T1 images were segmented, and surfaces were generated using FreeSurfer ^56^.

#### fMRI acquisition

fMRI data were acquired using a multi-echo T2*-weighted sequence. The sequence parameters were: TR = 1,057 ms, TEs = [14.2, 34.42, 54.64] ms, GRAPPA factor = 2, flip angle = 59°, matrix size = 80 x 80, slices = 52, multiband factor = 4, voxel size = 2.7mm isotropic.

#### Preprocessing

Functional MRI data was preprocessed using AFNI (afni_proc.py)^57^, incorporating alignment, distortion correction, multi-echo denoising, and regression modeling. For each participant, anatomical (T1-weighted) images were aligned to functional data, and all functional runs (n = 5) were concatenated (*tcat*). Multi-echo EPI data (three echoes; TE = 14.2, 34.42, 54.64 ms) were processed jointly, with motion correction parameters estimated from the second echo and applied to all echoes (*volreg*). Volumes were aligned to a minimum outlier reference and nonlinearly normalized to the MNI152 template (MNI152 2009c; *tlrc*). Distortion correction was performed using paired blip-up/blip-down fieldmap images.

Functional data were despiked to attenuate transient outliers and masked using an anatomical-functional intersection mask that was dilated (*mask*). Multi-echo data were optimally combined and denoised using multi-echo independent components analysis via tedana (*combine*) ^58–60^. The resulting time series were scaled to percent signal change (*scale*) and spatially smoothed with a 4 mm full-width at half-maximum Gaussian kernel (*blur*).

Nuisance regression (*regress*) included motion parameters and censoring of volumes exceeding motion (0.2 mm) or outlier (5%) thresholds. Task phases in which participants were speaking (**Fig. 1A**, Phase 2 and Phase 4) were explicitly modeled as regressors of no interest (dmBLOCK), accounting for variance associated with speech production without contributing to the residual time series used in subsequent analyses.

#### First-level modeling (trial-wise estimation)

Following preprocessing, single-trial responses were estimated using a general linear model (GLM) implemented via AFNI’s 3dDeconvolve. Analyses were performed on the residual time series from the preprocessing pipeline (errts), ensuring that nuisance effects (e.g., motion, baseline trends) had already been removed. Each image presentation was modeled as a separate regressor using a least-squares–separate approach (stim_times_IM with dmBLOCK), yielding one beta estimate per trial. The design matrix included one regressor per image presentation, allowing trial-level estimation of neural responses. Model parameters were estimated using AFNI’s 3dREMLfit to account for temporal autocorrelation. Resulting beta coefficients were retained for all trials and used in subsequent multivariate analyses.

#### Defining regions and networks of interest

The Schaefer parcellation ^29^ was used to designate 100 cortical regions. Parcels were assigned to large-scale functional networks according to the Yeo 17-network parcellation labels provided with the atlas ^30^.

### Computing neural shifts

Our goal was to detect within-subject changes in multivariate activity as participants viewed the same visual stimulus before and after being exposed to an alternative interpretation. Within each trial, we used an event-related general linear model (GLM) to estimate a beta weight at each voxel for each phase of interest (e.g., Initial Viewing and Second Viewing; **Fig. 1**). We then computed the cross-viewing similarity (Pearson correlation) between multivoxel activity patterns across phases within a given region (regions defined using the Schaefer parcellation). Correlation coefficients were Fisher z-transformed. To quantify neural change, we converted these values into “neural shift” scores by computing 1−*z*. Under this metric, pattern shift is minimal when two activity patterns are highly correlated and increases as the patterns become less similar.

### Statistical modeling of neural and behavioral measures

All analyses were conducted using linear mixed-effects models (LMEMs) implemented in lme4^61^. Parcel-level models included random effects for subject and image. Network-level models additionally included random effects for each parcel in the network. For models including eyetracking data, we also included a by-participant random slope for visual sampling dissimilarity, because this predictor varied within participant and was z-scored within participant. Thus, its fixed effect reflects within-subject variation in gaze change, with a one-unit increase corresponding to a one-SD increase relative to that participant’s own mean visual sampling shift. This random-effects structure follows recommendations that within-subject predictors include corresponding random slopes to avoid anti-conservative inference ^62^.

Semantic distance was modeled continuously in all main analyses. To allow for nonlinear effects, semantic distance was entered as a third-order orthogonal polynomial (poly(semantic_distance, 3)), which estimates linear, quadratic, and cubic components without collinearity. Unless otherwise specified, all statistics reported in the text reflect the linear component of this polynomial fit. For visualization, effects were generally plotted from companion models in which semantic distance was entered as a simple linear term to aid in the interpretability of the primary relationship of interest. References to effects at “high semantic distance” correspond to estimates evaluated at a semantic distance of 0.75; analogous estimates at low and medium semantic distance were evaluated at 0.25 and 0.50, respectively.

These values were used only to visualize and probe interactions in *post hoc* analyses, while the main inferential models treated semantic distance as continuous.

### Predicting neural shifts

#### Condition differences

Neural shift values were compared across conditions using LMEMs at both the parcel and network levels (**Fig. 2B**):

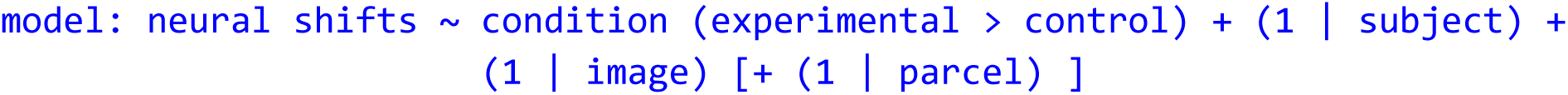

Where condition was contrast-coded (experimental > control). To account for the unequal number of trials across conditions (30 experimental vs. 15 control), we subsampled trials: for each participant, we randomly selected 15 experimental trials to match the control trial count and repeated this procedure 100 times. Models were fit separately for each subsampled dataset, and networks were considered significant if the condition effect reached significance in at least 99 of the subsamples.

### Predicting reinterpretation

To test whether reinterpretation was jointly predicted by semantic distance and neural shifts, we fit a linear mixed-effects model (**Fig. 3C**, **D**).

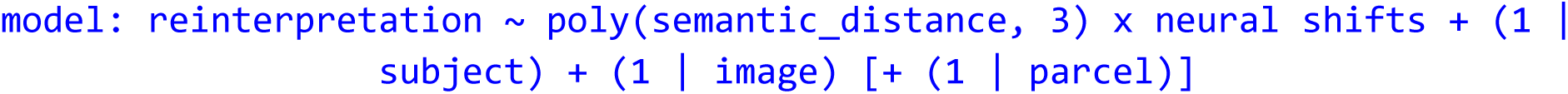

We then extended this network model to include both neural shifts and visual sampling shifts (operationalized as gaze allocation shifts or scanpath shifts; **Figs. 4, Supp Fig. 5D-G, Supp. Fig. 6**):

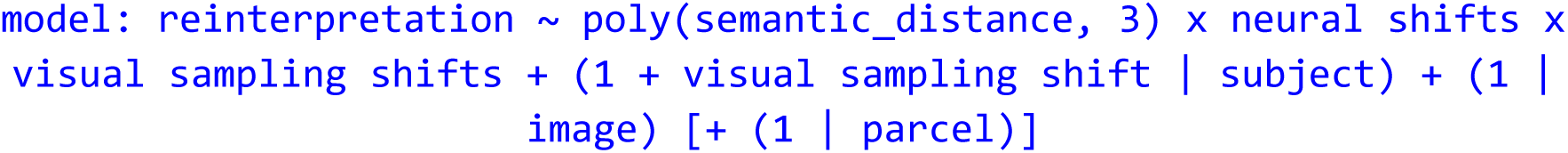

### Assessing shared variance between DMN-B and other association networks

To evaluate whether reinterpretation-related neural signals were independent across the higher-order association networks identified in our primary reinterpretation analysis (**Fig. 3C**: DMN-B, FPN-A, FPN-B, and VAN-B), or were instead reducible to variance shared with DMN-B, we fit a series of mutual adjustment models. For each network, neural shifts were averaged across parcels within that network for each subject and image, yielding a single trial-level neural shifts value per network per subject-image pair. We first fit an unadjusted model containing DMN-B neural shifts and their interaction with semantic distance. For each of the three comparison networks (FPN-A, FPN-B, VAN-B) we then fit two further models: an unadjusted model containing only that network’s neural shifts and their interaction with semantic distance, and a joint model containing both DMN-B and that network’s neural shifts, each interacting with semantic distance. The joint model supplies the adjusted estimate for both networks from a single fit. All models included random intercepts for subject and image, and semantic distance was entered linearly, so that coefficients are on a raw rather than orthogonal-polynomial scale; the same pattern of results holds when semantic distance is modelled with polynomial terms. From each model we extracted the neural shift x semantic distance interaction coefficient as the effect of interest.

DMN-B unadjusted model:

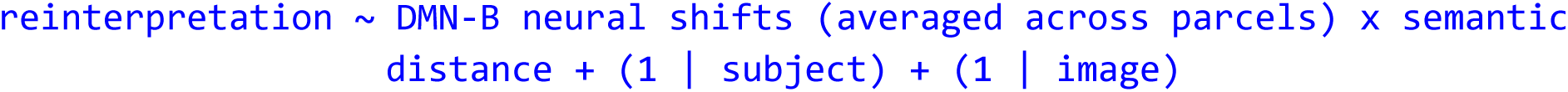

Comparison network unadjusted model (fit separately for each *Network* = FPN-A, FPN-B and VAN-B):

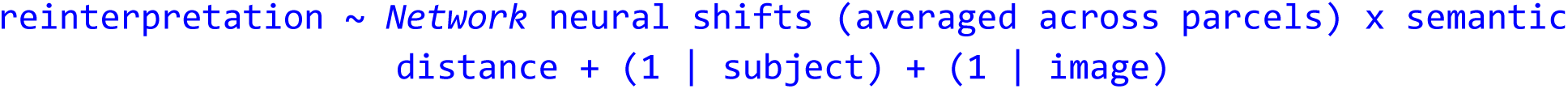

Joint adjusted model (fit separately for each *Network*, supplying the adjusted estimate for both network):

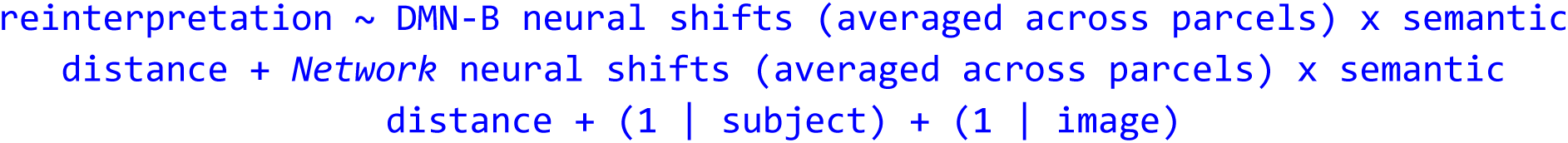

Stability of each network’s contribution to reinterpretation was assessed by comparing its neural shifts x semantic distance interaction coefficient before and after adjustment for DMN-B; a coefficient that remained stable in magnitude after adjustment was taken as evidence of an independent contribution, whereas a coefficient that markedly attenuated was taken as evidence that the network’s effect was largely reducible to shared variance with DMN-B. Note because neural shifts were averaged across parcels for this analysis, rather than including parcel as a random effect as in our primary parcel-level models, these coefficients are not directly comparable in scale to the whole-brain, parcel-level estimates.

### Comparing DMN-B and visual networks

To compare DMN-B and the visual networks directly, we fit a single model in which network (DMN-B or visual networks) was entered as a factor interacting with neural shift, visual sampling shift (gaze allocation or scanpath), and semantic distance.

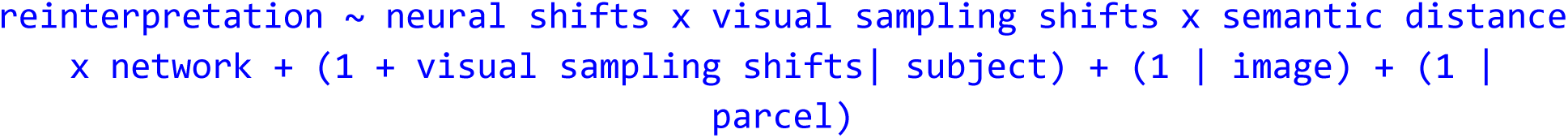

From this model, we estimated the marginal slope of neural shift on reinterpretation within each network at high semantic distance (0.75) and at low, mean, and high visual resampling shift (−1, 0, and +1 within-participant SD). The difference between the DMN-B and visual network slopes was then tested at each level of visual resampling shift. Marginal slopes and contrasts were computed with the *marginaleffects* package in R (slopes() and hypotheses() ^63^.

### Assessing shared variance between DMN-B and visual networks

To evaluate whether the reinterpretation-related neural signals observed in DMN-B and visual networks (VisCent/VisPeri networks combined in Fig. 4, independently in **Supplementary Fig. 6**) reflected unique or shared variance, we fit a series of mixed-effects models to trial-level data. Neural shifts were first averaged across parcels within each network to yield a single estimate per network per trial.

Each network was entered in the interaction structure through which its reinterpretation-related effect had emerged in the preceding analyses. DMN-B neural shift was modeled in interaction with semantic distance, because its relationship with reinterpretation did not significantly depend on visual resampling. Visual-network neural shift was modeled in a three-way interaction with semantic distance and visual sampling shift (either gaze allocation or scanpath), because its relationship with reinterpretation emerged specifically under conditions of substantial visual resampling. We fit three models.

The first estimated the unadjusted DMN-B effect:

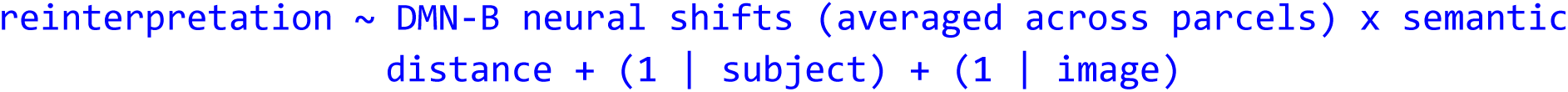

The second estimated the unadjusted visual-network effect:

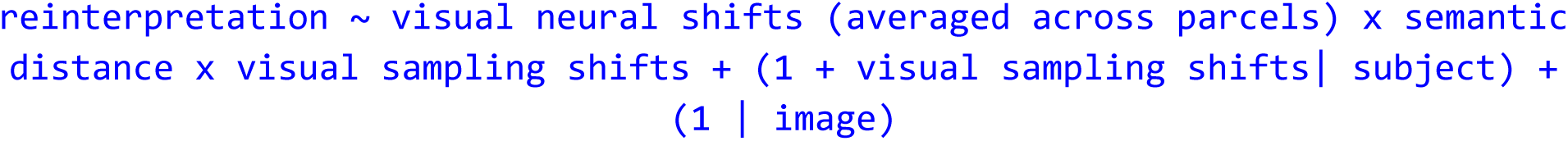

The third estimated the adjusted term by jointly including both networks.

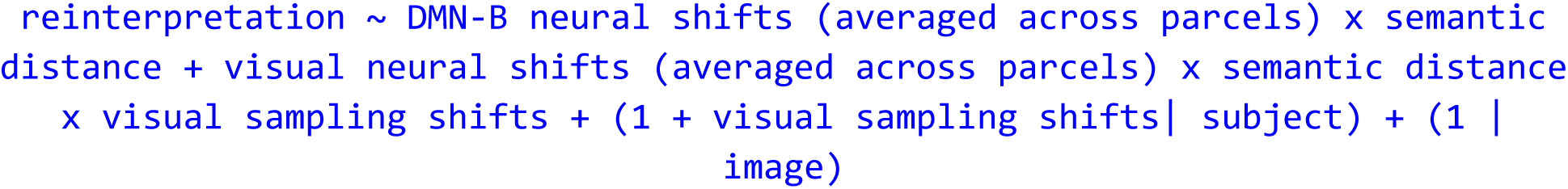

Stability of each network’s contribution was assessed by comparing the marginal slope relating its neural shift to reinterpretation, rather than the raw interaction coefficient used in the analyses above. Marginal slopes were used rather than individual interaction coefficients because DMN-B and the visual networks were modeled with different interaction structures: their raw coefficients therefore represented different model terms and were not directly comparable. Marginal slopes instead expressed both effects in a common quantity: the predicted change in reinterpretation associated with a one-unit increase in neural shift under specified conditions. Slopes were evaluated at high semantic distance (0.75) and, for the visual networks, at high visual sampling shift (+1 SD within subject). Marginal slopes were estimated via the delta method (*marginaleffects* package in R; ^63^).

### Network visualization and correction for multiple comparisons

For each analysis, false discovery rate (FDR) correction ^64^ was applied across the 17 networks to correct for multiple comparisons. Parcel-level results are shown uncorrected for visualization purposes. Network-level beta estimates and significance are reported in the corresponding figures. Cortical surface projections of parcel-level beta estimates were generated using SurfPlot ^65,66^. Predicted effects plots were generated using the ggeffects package in R ^67^. For visualization purposes, predicted effects are shown for the linear component only; full polynomial models are reported in the statistical analyses as noted above.

### Data and code availability

Analysis code is available at https://github.com/thefinnlab/mindlibs_rep. Raw MRI data will be deposited on OpenNeuro upon publication. The full stimulus set with annotations and the prompt scaffolds will be made publicly available upon publication.

### Use of Generative AI

During the preparation of this work, the author(s) used Claude (Anthropic) or ChatGPT (OpenAI) for assistance with programming data visualization code, syntax for analysis packages (e.g., marginaleffects, multimatch), and minor edits to the readability of the manuscript prose. All analyses were designed and conducted by the author(s), who reviewed and edited the AI-assisted code and take full responsibility for the content of the published article.

## Acknowledgements.

We thank Megan Leung, Sofia Yawand-Wossen, Evan Bloch, Georgia Nieh, Eli Bailit, David Igbalajobi, Isaac Wells, and Dallas Broderson for their help with fMRI data collection and/or early piloting of the behavioral paradigm. We also thank Clare Grall, Thomas Botch, Kathryn O’Nell, Zachary Paris, and especially Rekha Varrier, Qi Liang, Caroline Robertson and Monica Rosenberg for extensive discussions and feedback throughout the development of the paradigm, analyses and writing. Finally, we thank the Dartmouth Brain Imaging Center for their aid in data collection and feasibility, particularly Courtney Rogers and Terry Sackett.

Funding was provided by the NIH (R00MH120257 to E.S.F.; F31MH138084 to C.A.S.-S.) and by an NSF Graduate Research Fellowship (Award #2236868) to C.A.S.-S.

## Author Contributions

C.A.S.-S. and E.S.F. conceived the study. C.A.S.-S. programmed the task and analysis scripts. C.A.S.-S. and T.G.B. collected the data. C.A.S.-S. led the data analysis, with assistance from T.G.B. C.A.S.-S. and E.S.F. wrote the manuscript. All authors provided critical feedback, edited the manuscript and approved the final version.

