## Supplementary figures for "Shifts in neural representations, not gaze, track how people reinterpret ambiguous social scenes"

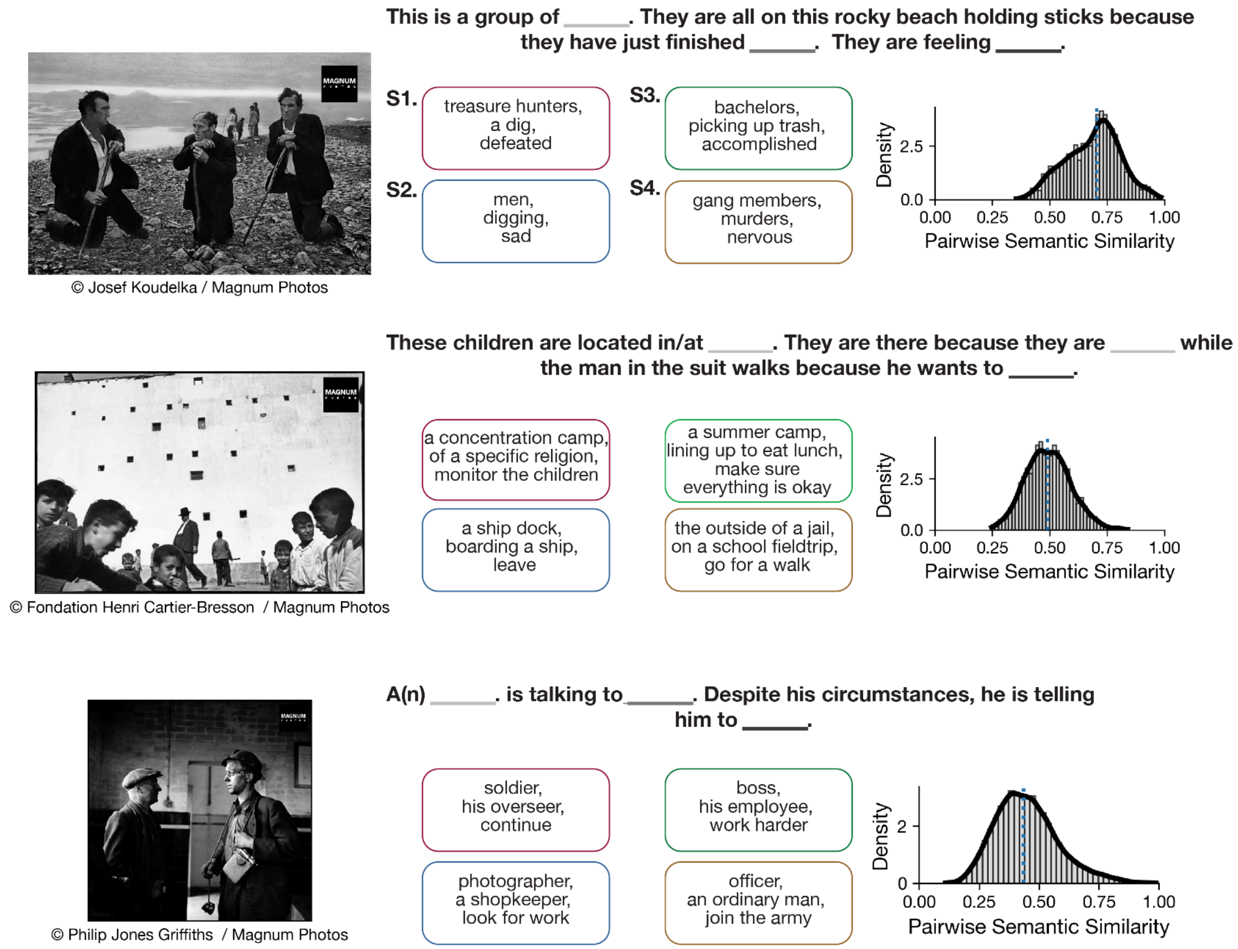


**Supplementary Fig. 1. Stimuli generate a wide range of interpretations.** Three example stimuli are shown with their shared scaffold, alongside interpretations from four representative subjects. Interpretations were encoded using miniLM-L6 1. Histograms show semantic similarity values between all possible pairs of subjects (N=56). See Methods section *Computing semantic distance* for details on the embedding procedure and how scaffold information was taken into account. The three example stimuli were photographs taken by Josef Koudelka (top), Henri Cartier-Bresson (middle) or Philip Jones Griffiths (bottom), sourced from Magnum Photos. The photographs used in the task were non-watermarked versions cropped to a uniform size (see *Methods* and Fig. 1B).

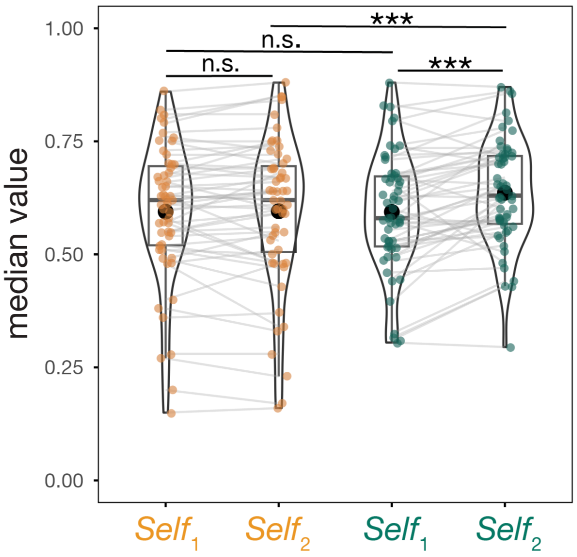


**Supplementary Fig. 2. Changes in self-ratings are specific to experimental trials.** Difference between $Self_1$ (initial confidence) and $Self_2$ (final self-rating) across experimental and control conditions (compare to **Fig. 2A**). Each dot represents a participant's median across trials (30 experimental, 15 control); gray lines connect the same participant across conditions. Statistics were derived from pairwise contrasts of estimated marginal means from a linear mixed-effects model predicting rating value from a condition × measure ($Self_1$/$Self_2$) interaction, with random intercepts for subjects and stimuli. $Self_1$ did not differ between conditions, nor did $Self_1$ and $Self_2$ differ within the control condition. $Self_2$ was significantly higher in the experimental than control condition, and $Self_1$ and $Self_2$ differed significantly within the experimental condition, with $Self_2$ higher than $Self_1$, indicating that updating was specific to experimental trials. Asterisks indicate significance ( *** $p < 0.001$).

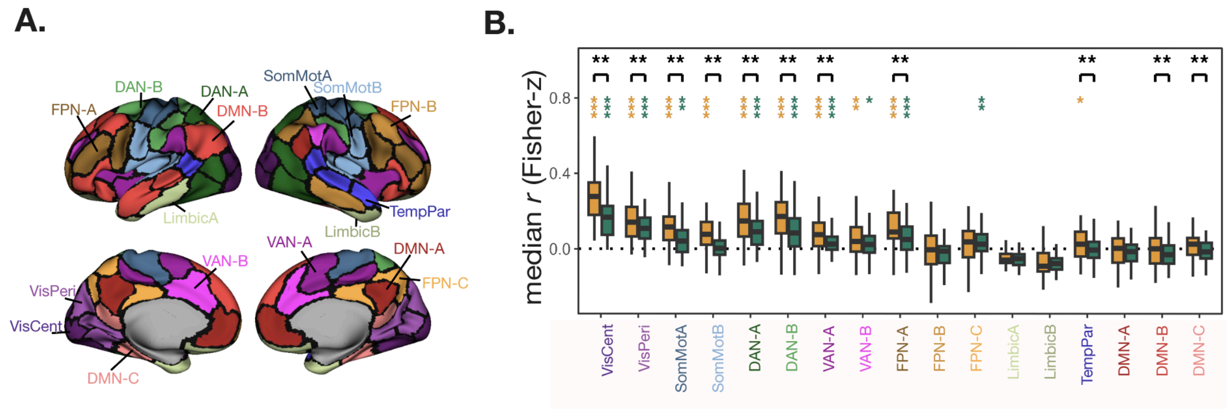


**Supplementary Fig. 3. Between-viewing similarity of neural representations decreases from sensory to association cortex.**

**A)** The 17 Yeo networks as assigned by the Schaefer parcellation [2,3], shown on an inflated cortical surface. Labels indicate a sample region within each network. Subnetworks belonging to the same larger network (e.g., DAN-A and DAN-B) are shown in different shades of the same color; parcels within the same subnetwork share the same color. Abbreviations: DAN, Dorsal Attention Network; VAN, Ventral Attention Network; FPN, Frontoparietal Network; DMN, Default Mode Network. **B)** Similarity between the initial and second viewing was quantified as the median Fisher *z*-transformed correlation across parcels within each network. Distributions represent subject-level median values separately for the control and experimental conditions. Similarity was significantly positive in the majority of networks and generally decreased from sensory to association cortex, with the weakest and least consistent effects in default mode regions. Colored asterisks indicate FDR-corrected one-sided one-sample tests of whether similarity was greater than zero within each condition (orange, control; green, experimental). Black asterisks above brackets indicate FDR-corrected experimental-versus-control contrasts from the linear mixed-effects model (see Fig. 2). Asterisks indicate significance (* $q < 0.05$, ** $q < 0.01$, *** $q < 0.001$).

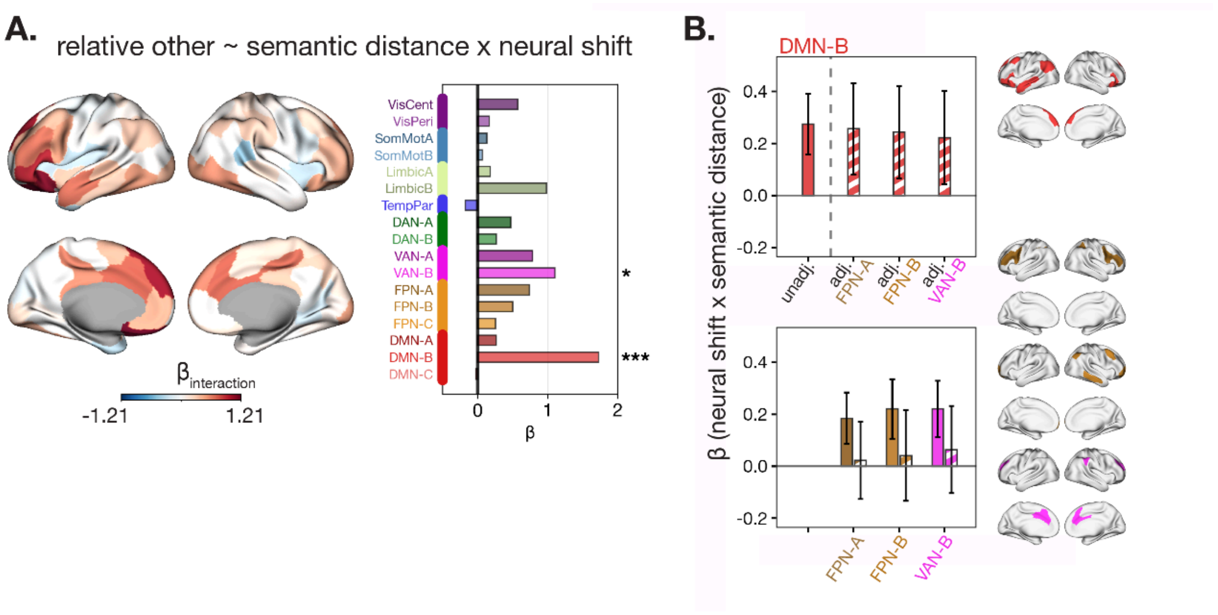


**Supplementary Fig. 4. A) The neural shift–reinterpretation relationship is robust to an alternative operationalization of reinterpretation.** Results of a linear mixed-effects model estimating reinterpretation — here computed as Other / (Self2 + Other), reflecting relative endorsement of the alternative — from the interaction between neural shifts and semantic distance, with random effects for parcel, subject, and stimulus. *Left.* Parcel-level β values projected onto the cortical surface. *Right*. Network-level summary of the interaction effect (β). Results are consistent with the primary operationalization (**Fig. 3C-D**), with significant effects in VAN-B and DMN-B. **B) DMN-B effects persist after adjustment for other association networks.** Bars show the β coefficient for the neural shift × semantic distance interaction from mutual adjustment models (see Methods, *Assessing shared variance between DMN-B and other association networks*). Top: DMN-B's effect unadjusted (solid, left of dashed line; β = 0.27) and after controlling for each comparison network (striped, right of dashed line; adjusted β = 0.26, 0.24, and 0.22 for FPN-A, FPN-B, and VAN-B, respectively). Bottom: Effects of FPN-A, FPN-B, and VAN-B unadjusted (solid; β = 0.18, 0.22, and 0.22) and after controlling for DMN-B (striped; adjusted β = 0.02, 0.04, and 0.06). DMN-B retained 81–94% of its unadjusted effect after adjustment, whereas the comparison networks retained 12–29%. Error bars reflect ±1 SE. Note that neural shifts were averaged across parcels within each network for this analysis, reducing power relative to the main analysis; results should be interpreted as a qualitative comparison of attenuation patterns rather than formal inference. Semantic distance was entered linearly here so that coefficients are on a raw rather than orthogonal-polynomial scale; the same attenuation pattern holds when semantic distance is modelled with polynomial terms.

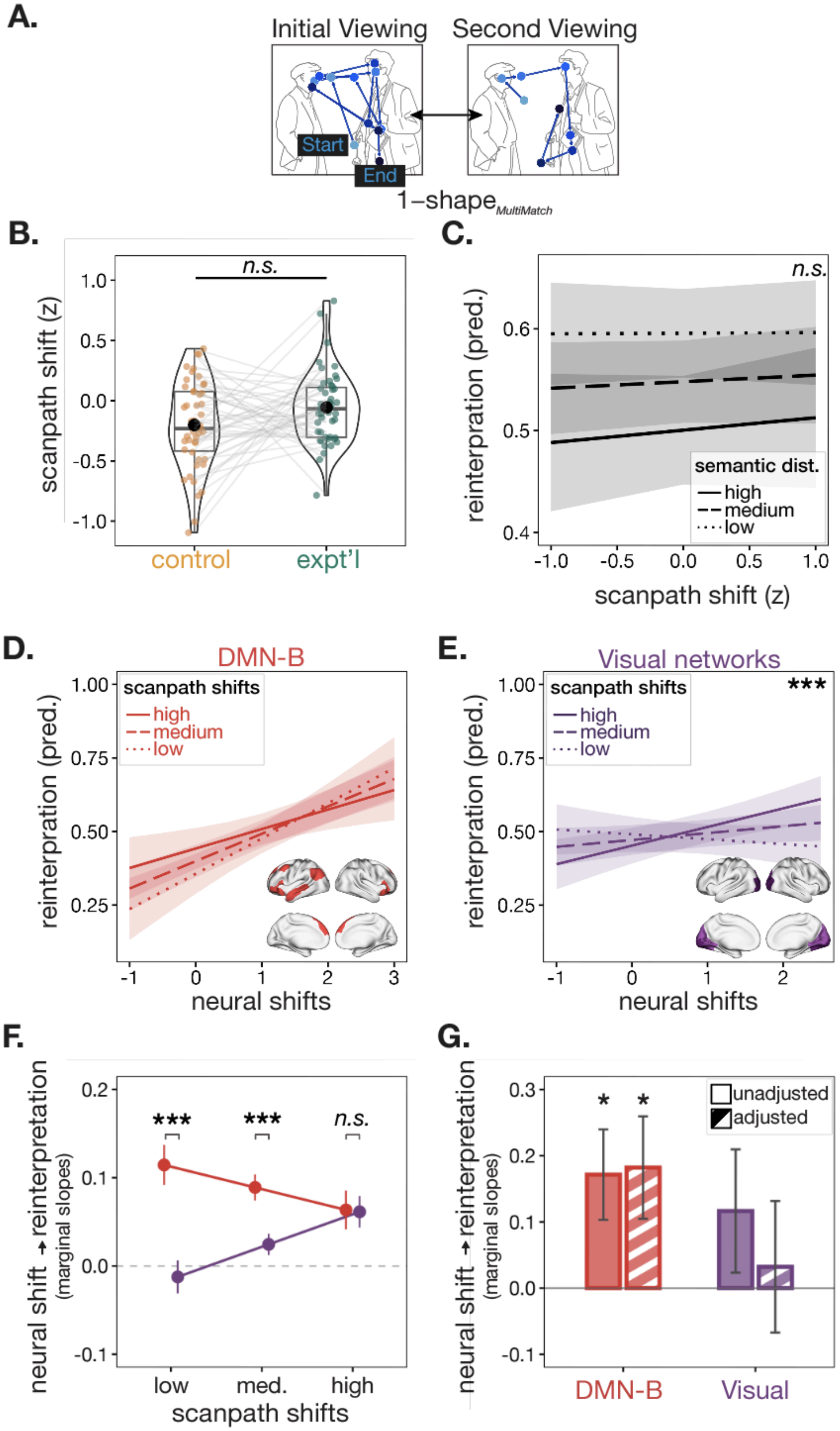
A.
Initial Viewing
Second Viewing
Start
End
1−shape MultiMatch
B.
n.s.
scanpath shift (z)
1.0
0.5
0.0
-0.5
-1.0
control
expt'l
C.
n.s.
reinterpration (pred.)
0.6
0.5
0.4
semantic dist.
high
medium
low
-1.0
-0.5
0.0
0.5
1.0
scanpath shift (z)
D.
DMN-B
scanpath shifts
high
medium
low
reinterpration (pred.)
1.00
0.75
0.50
0.25
-1
0
1
2
3
neural shifts
E.
Visual networks
scanpath shifts
high
medium
low
***
reinterpration (pred.)
1.00
0.75
0.50
0.25
-1
0
1
2
neural shifts
F.
neural shift → reinterpretation
(marginal slopes)
0.2
0.1
0.0
-0.1
***
***
n.s.
low
med.
high
scanpath shifts
G.
neural shift → reinterpretation
(marginal slopes)
0.3
0.2
0.1
0.0
-0.1
*
*
unadjusted
adjusted
DMN-B
Visual

**Supplementary Fig. 5. Visual sampling results replicate using scanpath shifts. A)** Sample scanpaths on a trial for the initial and second viewing. Scanpath shifts were computed by taking the shape metric from the Multimatch toolbox and converting it into a distance measure [4,5] (see Methods section *Computing scanpath shifts* for more detail). Color indicates time from first (dark) to last (light) fixation. Gaze data are from an actual trial, overlaid here on an author-created schematic representation of the original photographic stimulus for copyright purposes (see **Fig. 1**). **B)** Eyetracking results comparing scanpath shifts between conditions. Each dot represents the median across trials for a given participant. No significant differences were observed in how participants changed their sampling of the image between the initial and second viewing ($p > 0.05$; LMEM). **C)** Predicted fixed effects from a linear mixed-effects model estimating reinterpretation from the interaction between scanpath shifts and semantic distance. The interaction was not significant, indicating that at high semantic distance, scanpaths did not predict increased reinterpretation. **D)** Predicted fixed effects in the DMN-B network, highlighting that neural shifts predict reinterpretation at high semantic distance independently of scanpath shifts. **E)** Predicted effects in the visual networks, highlighting that neural shifts predict reinterpretation at high semantic distance specifically when scanpath shifts are large, consistent with a sensory role contingent on active visual resampling. **F)** Estimated marginal slope of neural shifts on reinterpretation (at high semantic distance) across low, medium, and high scanpath shifts, for DMN-B and the visual networks. DMN-B's slope is steeper than visual at low and moderate scanpath shifts, but not at high scanpath shifts. Error bars reflect ±1 SE. **G)** Marginal slope relating each network's mean neural shift to reinterpretation at high semantic distance and high scanpath shift, before (solid) and after (hatched) adjusting for the other network (see Methods, *Assessing Shared Variance Between DMN-B and visual networks*). Error bars reflect ±1 SE. Asterisks indicate significance (* $p < 0.05$, ** $p < 0.01$, *** $p < 0.001$).

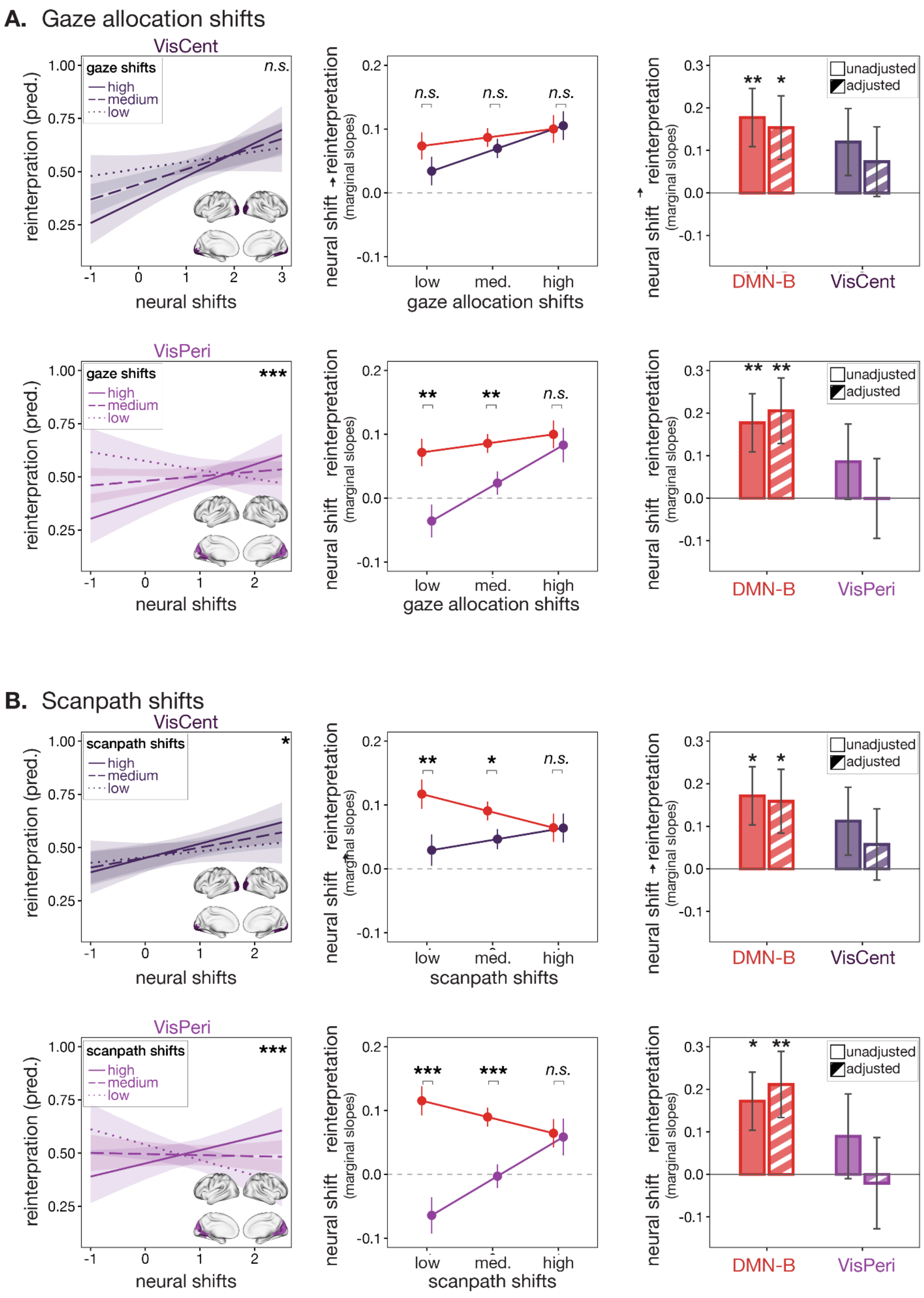


**Supplementary Fig. 6. Visual sampling results per visual network. A.** Results for VisCent (top row) and VisPeri (bottom row) from the gaze allocation shift metric (compare to main Fig. 4). **B.** Results for VisCent (top row) and VisPeri (bottom row) from the scanpath shift metric (compare to Supplementary Fig. 5). Asterisks indicate significance (* $p < 0.05$, ** $p < 0.01$, *** $p < 0.001$).
